# Single-cell analysis of human primary prostate cancer reveals the heterogeneity of tumor-associated epithelial cell states

**DOI:** 10.1101/2020.11.06.359802

**Authors:** Hanbing Song, Hannah N.W. Weinstein, Paul Allegakoen, Marc H. Wadsworth, Jamie Xie, Heiko Yang, Felix Y. Feng, Peter R. Carroll, Bruce Wang, Matthew R. Cooperberg, Alex K. Shalek, Franklin W. Huang

## Abstract

Prostate cancer is the second most common malignancy in men worldwide and consists of a mixture of tumor and non-tumor cell types. To characterize the prostate cancer tumor microenvironment, we performed single-cell RNA-sequencing on prostate biopsies, prostatectomy specimens, and patient-derived organoids from localized prostate cancer patients. We identify a population of tumor-associated club cells that may act as progenitor cells and uncover heterogeneous cellular states in prostate epithelial cells marked by high androgen signaling states that are enriched in prostate cancer. *ERG*- tumor cells, compared to *ERG*+ cells, demonstrate shared heterogeneity with surrounding luminal epithelial cells and appear to give rise to common tumor microenvironment responses. Finally, we show that prostate epithelial organoids recapitulate tumor-associated epithelial cell states and are enriched with distinct cell types and states from their parent tissues. Our results provide diagnostically relevant insights and advance our understanding of the cellular states associated with prostate carcinogenesis.

## Introduction

The prostate consists of multiple cell types including epithelial, stromal, and immune cells, each of which has a specialized gene expression profile. The development of cancer from prostate tissue involves complex interactions of tumor cells with surrounding epithelial and stromal cells and can occur multifocally, suggesting that prostate epithelial cells may undergo cellular state transitions towards carcinogenesis^1–6^. Previous studies on prostate cancer (PCa) molecular changes have focused on unsorted bulk tissue samples, leaving a gap in our understanding of the adjacent epithelial cell states.

The classification of prostate epithelial cells has been expanded over the past few years from three types (basal epithelial cells, luminal epithelial cells, and neuroendocrine)^7, 8^ to include hillock cells and club cells^9^. The roles of these additional cell types in the prostate are largely unknown. Most PCa are marked by the expansion of malignant cells with luminal epithelial features and the absence of basal epithelial cells. However, to date, the role of additional cell populations beyond the luminal and basal types is not well known.

Another underexplored area is the tumor microenvironment changes that occur based on dominant genomic drivers in PCa. PCa tumor cells are driven by a number of oncogenic alterations that include highly prevalent gene fusion events involving ETS family transcription factors, such as *TMPRSS2*-*ERG* and *ETV1*/4/5^1, 10–12^. Tumor cells without ETS fusion events and non-malignant luminal cells, however, have not been thoroughly characterized on a single-cell level, and uncertainty remains whether ETS fusion events could evoke differential stromal and immune cell responses.

To characterize tumor cells and the surrounding epithelial, stromal, and immune cell microenvironment and identify cell states that are associated with tumorigenesis, we performed single-cell RNA-sequencing (scRNA-seq) on PCa samples. Furthermore, we derived *in vitro* organoids from PCa tumor tissues followed by scRNA-seq to chart molecular and cellular changes in prostate cancers from localized PCa patients. We aimed to understand at single-cell resolution the tumor microenvironment and cellular states associated with prostate carcinogenesis.

## Results

To probe the diversity of cell types and transcriptional states of cells in localized prostate cancer specimens, we isolated single cells from biopsies and surgical resection specimens from men with localized prostate cancer for scRNA-seq (**Supplemental Table 1)** using an improved Seq-well single-cell platform^13^. Altogether, 21,743 cells were analyzed and a total of nine different major cell types were identified, marked by specific gene expression profiles (**Methods**, **Figure 1a,b**).

**Figure 1.**
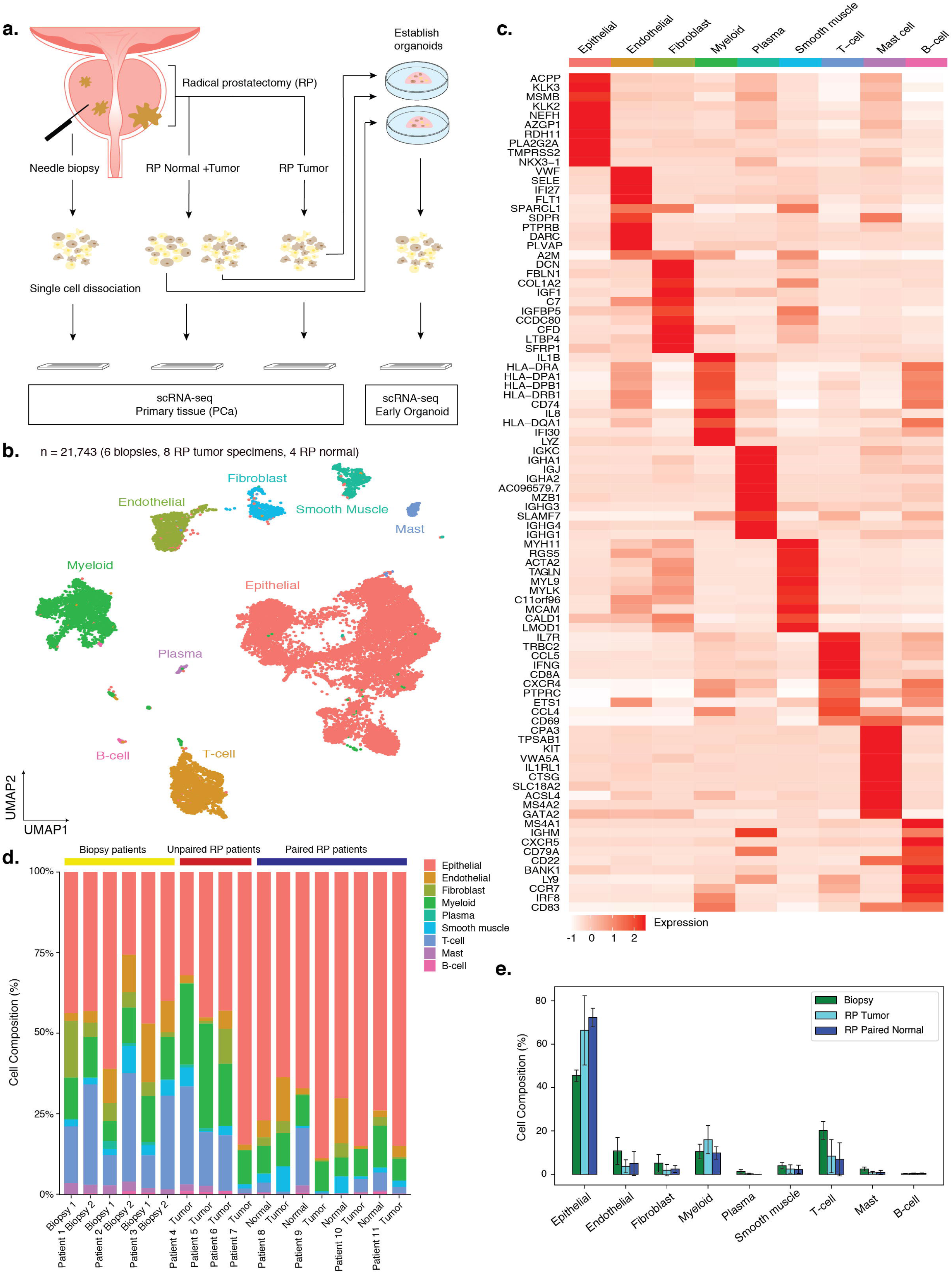
PCa sample single-cell RNA-sequencing overview and identification of major cell types in localized prostate cancer. **a.** Single-cell RNA-sequencing workflow on PCa biopsies, radical prostatectomy (RP) specimens, and *in vitro* organoid cultures grown from RP tumor specimens using the Seq-Well platform. **b**. Overview of major cell types identified within the combined dataset consisting of 21,743 cells from all biopsies (N = 6) and RP specimens (N = 12). Cell types are labeled in colors from corresponding clusters in the UMAP. **c.** Heatmap for the top 10 differentially expressed genes in each cell type. **d.** Cell type composition stacked bar chart by sample. Cell counts for each sample are normalized to 100%. Sample type is annotated (top) and patients are labeled below the x-axis. **e.** Cell composition comparison for each cell type among three sample types: biopsy patients (N = 3), RP tumor specimens (N = 8), and RP paired normal tissues (N = 4). Mean and confidence interval for each cell type are indicated in the grouped bar chart.

Cell type identification was determined by examining differentially expressed genes (DEGs) as well as signature scores from normal prostate and immune cell population gene sets^9, 14^. Cells in the merged dataset were annotated as epithelial, stromal (endothelial, fibroblast, and smooth muscle), and immune (T-cells, myeloid cells, plasma cells, mast cells, and B-cells) cells based on established marker genes. Epithelial cells (N = 13,322) were identified based upon the expression of luminal epithelial (LE) markers *KLK3, ACPP, and MSMB,* consistent with LE cells found as the dominant epithelial cell type in PCa samples. Immune cells were identified based on the high-level expression of *PTPRC* in five clusters, of which one cluster was marked by high-level expression of *IL7R*, *CD8A,* and *CD69,* indicating a mixture of both CD8 and CD4 T-cells; a second cluster was characterized by the myeloid cell markers *APOE, LYZ,* and *IL1B*^15–18^. The third *PTPRC+* cluster represented plasma cells marked by high level expression of *MZB1* and *IGJ.* The other two remaining *PTPRC+* clusters were annotated as mast cells expressing *CPA3, KIT,* and *TPSAB1*, and a population of B-cells expressing *MS4A1, CD22,* and *CD79A*. Stromal cells in our dataset consisted of endothelial cells characterized by *CLDN5* and *SELE* expression, fibroblasts expressing *C1S, DCN,* and *C7*, and smooth muscle cells expressing *ACTA2*, *MYH11,* and *RGS5* (**Figure 1c**).

As our samples consisted of prostate biopsies (N = 3 patients) and radical prostatectomy (RP) specimens (N = 8 patients), half of which had matched benign-appearing tissue (**Supplemental Table 1**), we tested whether each sampling strategy captured a similar distribution of different cell types across samples. All major cell types were captured in each sample with epithelial cells comprising the largest population (**Figure 1d**). No significant difference was found among the three sample types (p > 0.05, Mann Whitney U-test) (**Figure 1e**). We also compared the cell type composition among paired tumor (N = 4), paired normal (N = 4), and RP unpaired tumor tissues (N = 4) (**Supplemental Table 1**) and found no significant differences (p > 0.05, Mann-Whitney U test). The main cell types identified were validated by SingleR annotation^19^ (**Supplemental Table 1**). Furthermore, within each biopsied patient, we tested whether biopsies from the two anatomical regions identified similar cell types and found that all cell types were recovered in each biopsy sample with some sampling differences by anatomical regions **(Supplemental Table 1).**

### Epithelial cell clusters reveal tumor cells and non-tumor surrounding epithelial cell heterogeneity

To identify the transcriptional cell states of epithelial cells associated with prostate cancer, we performed a graph-based clustering analysis and identified 20 clusters (**Figure 2a**). We then conducted single-sample gene set enrichment analyses^20, 21^ (ssGSEA) using signature gene sets developed previously from single-cell profiling of normal prostates **(Supplemental Table 2)** to determine the major cell subtypes^9^. Clusters with *KRT5*, *KRT15*, *KRT17,* and *TP63* expression (**Figure 2b**) and significantly upregulated basal epithelial (BE) signature scores were identified as BE cells. Given that tumor cells predominantly express LE cell markers such as *KLK2, KLK3, ACPP,* and *NKX3-1*, clusters with high LE signature scores could be either tumor cells or non-malignant LE cells (**Figure 2b**). BE and LE signature feature plots also revealed a cluster of cells (cluster 5) that we termed other epithelial (OE) cells (**Figure 2a,c**), with lower BE and LE signatures scores (**Supplemental Figure 1a**), and were characterized by *PIGR, MMP7,* and *CP* expression. In previous studies, *PIGR* has shown a role in promoting cell transformation and proliferation^22^; *MMP7* may promote prostate carcinogenesis through induction of epithelial-to-mesenchymal transition^23^, and serum *CP* levels have been used as a marker in PCa^24^ (**Figure 2b)**.

**Figure 2.**
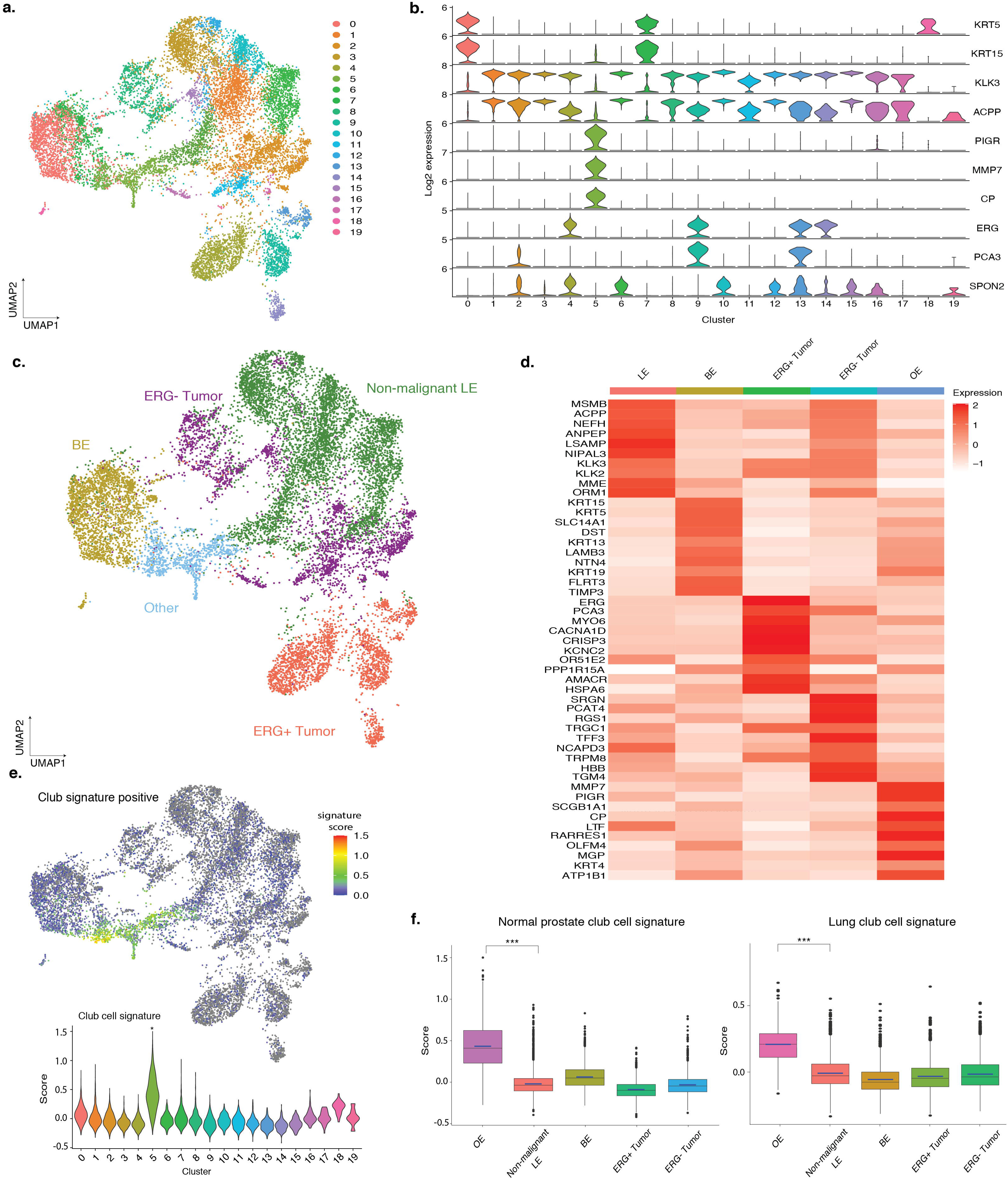
Identification of tumor cells and major epithelial cell types including club cells. **a**. UMAP projection of all 20 clusters identified in the epithelial cells. Clusters are labeled in the UMAP. **b.** Violin plots of representative marker genes across the clusters. **c.** UMAP of epithelial cells annotated by cell types. **d.** Heatmap for the top 10 differentially expressed genes in each cell type. **e.** Club cell signature scores of each epithelial cell projected on the UMAP and signature score violin plots across all clusters. **f.** Box plots of club cell signature scores from normal club cells and lung club cells across epithelial cell types (***: p < 0.001, Wilcoxon rank sum test).

Approximately 50% of PCa tumors from European ancestry patients harbor *TMPRSS2-ERG* fusion events and less frequently harbor other ETS fusion events (*ETV1, ETV4, ETV5*)^25^. To identify tumors cells, we tested cells for expression of *ERG, ETV1, ETV4,* and *ETV5*. *ERG* expression was found upregulated in four clusters (**Figure 2b, Supplemental Figure 1a**); therefore, we annotated these four clusters as *ERG*+ tumor cells that comprised cells from six patients. Other than tumor cell clusters, only endothelial cells showed high-level *ERG* expression. The identity of *ERG*+ tumor cells was further supported by the upregulation of the SETLUR PROSTATE CANCER TMPRSS2 ERG FUSION UP gene set signature score in these cells^26^. Furthermore, STAR-Fusion^27^ identified potential fusion transcripts of *TMPRSS2-ERG* fusion in two *ERG*+ patients. No clusters with *ETV1, ETV4,* or *ETV5* expression were detected (**Supplemental Figure 1b**).

To identify tumor cells without ETS fusion events, we tested the LIU PROSTATE CANCER UP and other PCa tumor marker gene set signature scores and identified seven clusters in total with upregulated signature scores of at least one prostate cancer gene set (**Supplemental Figure 1a,b**). Single sample gene set enrichment analysis (ssGSEA) on these 11 clusters also showed at least one prostate cancer gene set that scored in the top 1% of all C2 CGP gene set collection (N = 3,297) (**Supplemental Table 3**). Therefore, we classified four clusters with *ERG* expression as *ERG*+ tumor cell clusters and the other seven as *ERG*- tumor cells (**Figure 2c**). All *ERG*- tumor cell clusters expressed tumor marker *SPON2*^28^ (**Figure 2b**).

To validate our tumor cell assignments, we estimated copy number variants (CNV) via InferCNV^29^, using non-malignant LE cells as a reference. From the CNV estimation visualization (**Supplemental Figure 1c**), we identified significantly different CNV patterns in both *ERG*+ and *ERG*- tumor cells. Non-uniform CNV profiles were detected within *ERG+* and *ERG-* tumor cell populations, suggesting heterogeneity in both tumor cell populations.

While we did not observe a separate neuroendocrine cell cluster, we tested for prostate neuroendocrine (NE) cells^9, 30^ using an established NE cell signature gene set^9^ and computed the NE signature scores for each epithelial cell. Taking the cells ranking in the top 0.5% NE signature score, we detected 66 putative NE cells within the BE cell population, characterized by *CHGB*, *KRT4,* and *LY6D* expression^9^ (**Supplemental Figure 1c,d**).

To examine if our annotation method could accurately identify each cell type, we computed the top 10 biomarkers for each cell type (**Figure 2d**). BE cells showed high expression of established basal epithelial cell markers *KRT5, KRT15,* and *KRT17*. The top biomarkers in the OE clusters were *PSCA, PIGR, MMP7*, *SCGB1A1, and LTF*, of which *PSCA* is considered to be a prostate progenitor cell marker enriched in PCa^31–33^ and *SCGB1A1* a marker for lung club cells^34^. *ERG*+ and *ERG*- tumor cells and non-malignant LE cells all showed high expression of luminal markers *KLK3, KLK2, and ACPP*^35^. *ERG*+ tumor cells were characterized by expression of *ERG* and tumor markers *PCA3, AMAC,* and *TRPM8*^35–37;^ *ERG-* tumor cells were marked by the expression of tumor markers *PCA3* and *TRPM8*^35–37^ (**Figure 2d**).

Since most PCa are androgen-responsive with tumor cell proliferation dependent on the activity of the androgen receptor (*AR*)^36–39^, we tested for androgen responsiveness among the epithelial cell populations and identified LE cells and tumor cells as the most androgen responsive due to significantly higher *AR* signature scores compared to the other epithelial cell types (**Supplemental Figure 1a**). To identify putative prostate cancer stem cells that may contribute to prostate cancer development, we used an adult stem cell signature gene set^38^ and found that 56.4% of the BE cell population was enriched for the stem cell signature (**Supplemental Figure 1b**).

A previous single-cell study of normal human prostate reported two populations of other epithelial cells: hillock cells characterized by *KRT13, SERPINB1, CLDN4,* and *APOBEC2* expression and club cells characterized by the expression of *SCGB3A1*, *PIGR*, *MMP7*, *CP,* and *LCN2*^9^. While we did not detect a separate hillock cell population within our prostate cancer epithelial cells (**Supplemental Figure 1e**), we did detect a distinct population representing 6.5% of all epithelial cells (872 of 13,322) characterized by expression of *PIGR, MMP7, CP,* and *LTF* (**Figure 2d**) (FDR q < 10e-20). We hypothesized that this epithelial cluster represented club cells that had previously been described in lung^34^ and normal prostate specimens^9^. To test this hypothesis, we applied a normal prostate club cell signature gene set^9^ and projected the signature onto our epithelial UMAP. We found that cells with high club cell signature scores largely overlapped with this OE cluster (cluster 5) (**Figure 2e**). Furthermore, this cluster was enriched for a lung club cell signature compared to other clusters (p < 0.001, Wilcoxon rank sum test) (**Figure 2f**). Based on these results, we annotated this cluster as club cells. We then conducted an ssGSEA analysis on all epithelial cells using the BE, LE, and club cell signatures generated from the DEG profiles (**Supplemental Table 2**). All three cell type signature scores were strongly correlated to the corresponding cell types, supporting our annotation (**Supplemental Figure 1f**).

### Club and BE cells harbor PCa-enriched LE-like cell states that are upregulated in *AR* signaling

A recent study identified a luminal progenitor cell type in mouse and human prostates characterized by high expression of LE markers *KRT8*, *KRT18,* and other markers including *PSCA, KRT4*, *TACSTD2,* and *PIGR*^39^. In both normal and PCa epithelial cells datasets in our study, we could not identify a single cell type distinguished by the co-expression of *KRT8*, *KRT18,* and *TACSTD2*; however, *PSCA* and *PIGR* were expressed at higher levels in club cells compared to other epithelial cell types (**Supplemental Figure 2a**), indicating that the luminal progenitor cells previously identified are most similar to the club cells in our analysis.

Club cells in PCa have not been previously characterized. Since we exclusively captured club cells but not hillock cells in our PCa samples, we hypothesized that club cells play a role in carcinogenesis. To test this hypothesis, we integrated our prostate cancer club cells (Club PCa) with normal club cells from a previous study from healthy controls^9^ (Club Normal) and detected six cell states with distinct transcriptomic profiles (**Figure 3a**) by selecting an optimal resolution to yield stable clusters (**Supplemental Figure 2b**). Overall, compared to club cells from normal samples, PCa club cells exhibited downregulation of genes including lipocalin 2 (*LCN2*) and growth-inhibitory cytokine *SCGB3A1*^40, 41^ and upregulation of *LTF, AR,* and *AR* downstream members including *KLK3, KLK2, ACPP,* and *NKX3-1* (**Figure 3b**), which we hypothesized could be driven by the enrichment of one or more specific club cell states in the PCa samples.

**Figure 3.**
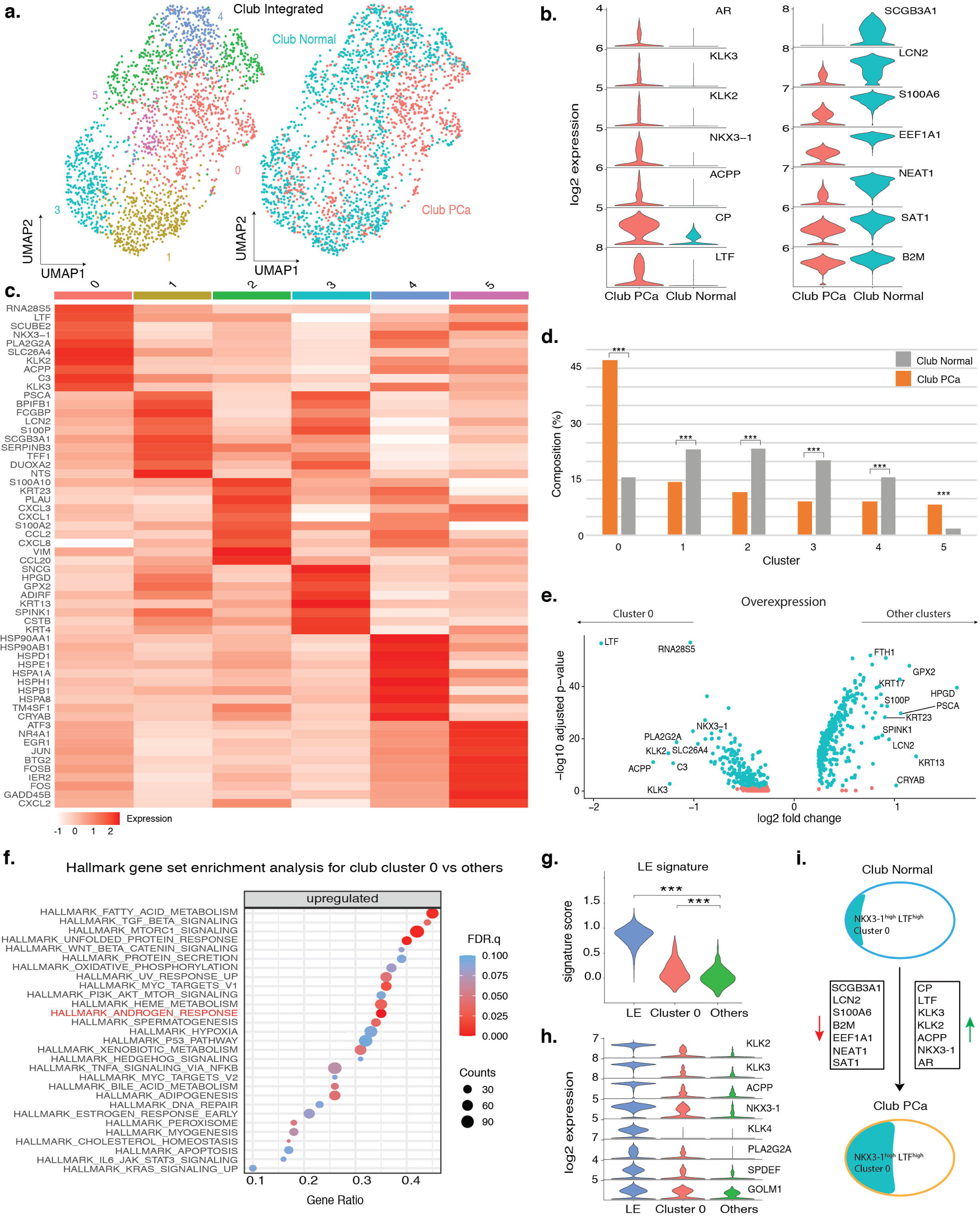
Identification of PCa-enriched club cell states with upregulated androgen response signature. **a.** UMAP of integrated club cells from PCa samples (Club PCa) and club cells from normal samples (Club Normal), color coded by cell states with differential gene expression profiles (left) and sample type (right). **b.** Violin plots of representative marker genes between the two types of club cells. **c.** Heatmap for the top 10 differentially expressed genes in each cell state. **d.** Grouped bar chart comparison of 6 cell state compositions between Club PCa and Club Normal. Significance levels are labeled (***: FDR q < 0.001, Wilcoxon rank sum test). **e.** Volcano plots of the overexpressed genes in Club cell cluster 0 and other cell states within the PCa samples. **f.** Top 20 upregulated signaling pathways between Club cell cluster 0 and the other club cells on Hallmark gene set collection (N = 50) within the PCa samples. Gene counts for the corresponding gene set indicated by marker radius. Statistical significance levels (FDR) are shown by color gradient. **g.** Comparison of LE signature scores between Club cluster 0 and other club cells (***: p < 0.001, Wilcoxon rank sum test) within the PCa samples. **h.** Violin plot comparison between Club cluster 0, other club cells and LE for multiple LE markers within the PCa samples. **i.** Schematic marker of gene expression changes between Club Normal and Club PCa. Gene downregulation and upregulation in Club PCa compared to Club Normal represented by red and green arrows. Proportion of Club cell cluster 0 within all club cells represented by area in blue and characterized by its LE-like state and high-level expression of *LTF* and *NKX3-1*.

For each of the six subclusters, a group of distinctive DEGs was identified (**Figure 3c**) and each subcluster was detected in both Club PCa and Club Normal (**Supplemental Figure 2c**), of which, Club PCa was significantly enriched in cluster 0 by more than three-fold compared to Club Normal (p < 0.001, Fisher’s exact test (FET)) (**Figure 3d**). This cluster was distinguished by a higher level of expression of *LTF*, luminal markers, and downstream *AR* pathway molecules *KLK2*, *KLK3*, *ACPP, PLA2G2A,* and *NKX3-1* (**Figure 3e**), suggesting a luminal-like and androgen-responsive state^39^.

To test the functional role of this cell state, we performed GSEA analysis using C2 canonical pathways (N = 2,232) (**Supplemental Table 4)** and Hallmark (N = 50) gene set collections on cluster 0 vs other cell states. Among the top significantly upregulated gene sets in cluster 0 was the Hallmark Androgen Response pathway (FDR q < 10e-5) (**Figure 3f**). These results were consistent with the upregulation of downstream *AR* pathway molecules in cluster 0.

Next, we tested whether this PCa-enriched cell state represented a luminal-like cell state. We observed higher LE signature scores in cluster 0 compared to other cell states (p < 0.001, Wilcoxon rank sum test) (**Figure 3g**). Specifically, we compared the expression levels of all LE markers among cluster 0, other club cells, and the LE population within the PCa samples, and found that Club cell cluster 0 exhibited higher expression of *KLK2, KLK3, ACPP, NKX3-1, KLK4, PLA2G2A, SPDEF* and *GOLM1* than other club cells (**Figure 3h**). While *AR* itself was not upregulated in cluster 0 (**Supplemental Figure 2d**), *KLK4,* a regulator of androgen response signaling, was upregulated in this cell cluster^42^.

Overall, the population of PCa club cells, compared to normal prostate clubs, was characterized by higher androgen signaling and an enrichment of an *LTF^high^* and *NKX3-1^high^* luminal-like cell state (**Figure 3i**).

The finding of a luminal-like club cell state led us to investigate if a similar cell state existed in the BE cell population of prostate cancer samples. Therefore, we integrated BE cells in the PCa samples (BE PCa) with BE cells from normal samples (BE Normal) and identified nine cell states (**Figure 4a**, **Supplemental Figure 3a)** with distinctive DEGs (**Supplemental Figure 3b**). While all nine cell states were represented in both BE PCa and BE Normal cells (**Figure 4b**), BE PCa was found to be significantly enriched in cluster 6 (31.8% vs 0.2%, PCa vs Normal) while BE Normal was enriched in cluster 4 (0.8% vs 15.9%, PCa vs Normal) (FDR q < 10e-5, FET; **Figure 4b**, **Supplemental Figure 3c**). This BE cluster 6 was marked by higher expression of downstream *AR* pathway members *KLK3, KLK2,* and *ACPP* **(Supplemental Figure 3b**). Compared to other BE cells in the PCa samples, BE cluster 6 also showed significant upregulation of *AR* (p < 0.01, Wilcoxon rank sum test, **Supplemental Figure 3d**). Among the top significantly upregulated gene sets were the Hallmark Androgen Response pathway within the Hallmark gene set collection **(Supplemental Table 4**), as well as androgen response pathways, estrogen pathways, the insulin signaling pathway, and Kegg pathways in cancer within the C2 CP gene set collection (FDR q < 0.1, Wilcoxcon rank sum test) (**Figure 4d**)^42–44^. As *AR* pathway members were among the top biomarkers for cluster 6 (**Figure 4e**), we hypothesized that BE cluster 6 may represent an intermediate BE/LE cell state, even though it did not cluster separately from other BE cells on the epithelial cell UMAP (**Figure 4f**). Therefore, we compared the expression levels of LE markers in BE cluster 6 and found that BE cluster 6 was upregulated in multiple LE markers compared to other BE cell states (**Figure 4g**), though at lower levels compared to the PCa LE cell population. Moreover, we found that BE cluster 6 was significantly upregulated in the Hallmark Androgen Response signature (p < 0.001, Wilcoxon rank sum test) and LE signature score (**Figure 4h**), supporting that this cell state may be a luminal-like state associated with prostate cancer.

**Figure 4.**
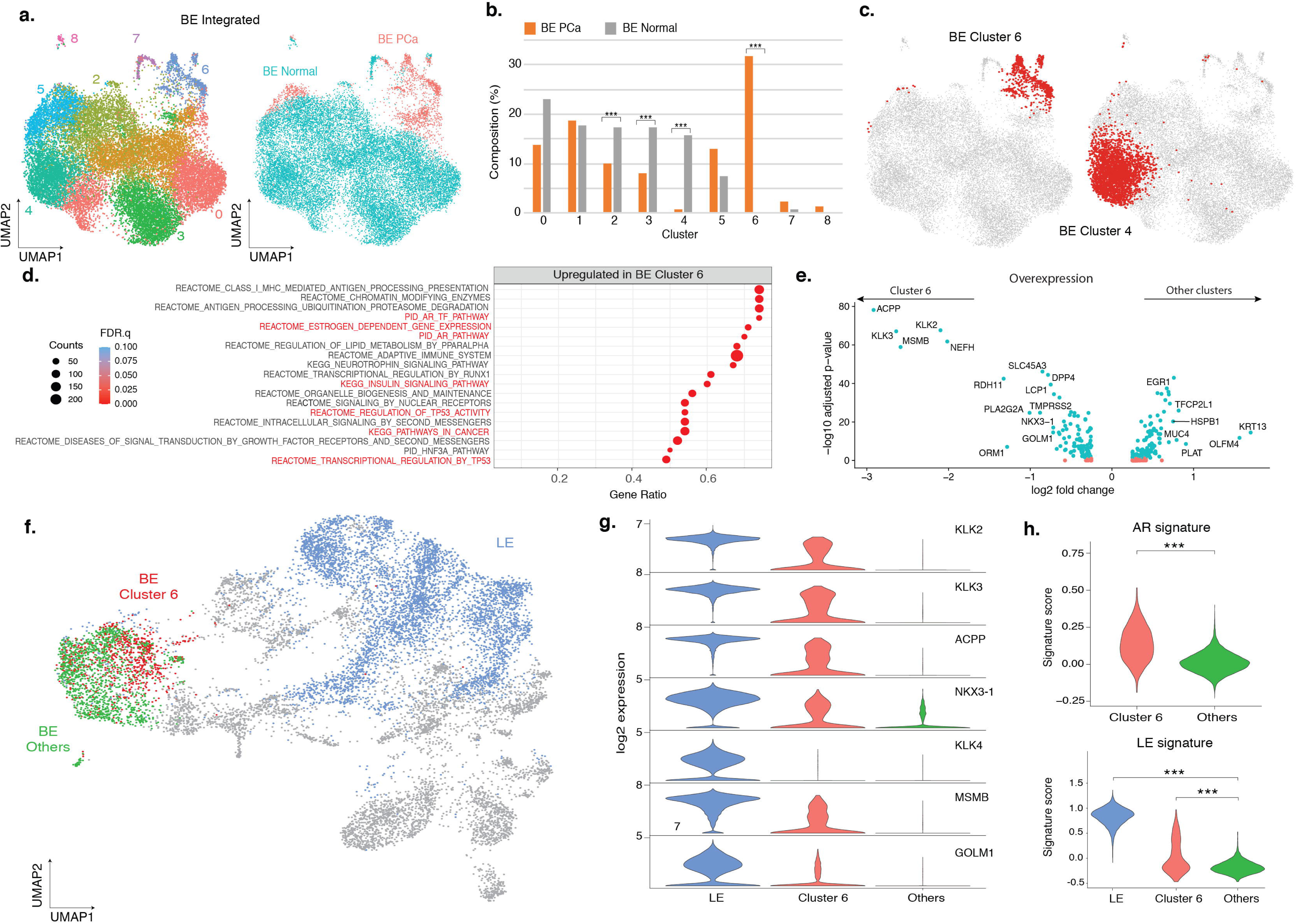
Integration of BE and LE cells identifies tumor-associated cell states enriched in the PCa samples. **a.** UMAP of integrated BE cells labeled by cell states (left) or samples type (BE PCa and BE Normal) (right). **b.** Cell composition comparison between BE PCa and BE Normal. **c.** PCa and normal enriched cell states 4 and 6 highlighted in the integrated BE UMAP. **d.** Top 20 upregulated signaling pathways between cluster 6 and the other BE on C2 canonical gene set (C2CP) collection (N = 2,332). Gene counts for the corresponding gene set are indicated by marker radius. Statistical significance levels (FDR) are shown by color gradient. Pathways associated with PCa tumor progression and invasiveness are highlighted in red. **e.** Volcano plots of the overexpressed genes in BE cluster 6 and other BE cell states within the PCa samples. **f.** Distribution of BE cluster 6, other BE and LE on the overall epithelial cell UMAP. **g.** Violin plot comparison between BE cluster 6, other BE and LE for multiple LE markers within the PCa samples. **h.** Comparison of Hallmark *AR* pathway signature and LE signature scores within the PCa samples (***: p < 0.001, Wilcoxon rank sum test).

Similarly, we identified eight cell states within the integrated LE dataset (**Supplemental Figure 3e**). Unlike BE and club cells, we observed a clear separation between LE PCa and LE Normal (**Supplemental Figure 3e**). LE PCa was significantly enriched in four cell states and LE Normal significantly enriched in two (p < 0.001, FET) (**Supplemental Figure 3f**). Cluster 5 was marked by co-expression of club cell markers such as *PIGR, MMP7* and *CP*, suggesting an intermediate population of LE and club cells. Cluster 1 was characterized by the overexpression of the *AR*-regulated gene *TMEFF2* and insulin-like growth factor *IGFBP5* compared to other cell states, and cluster 2 was upregulated in *AR* expression (**Supplemental Figure 3g**).

We then tested if the PCa-enriched cell states in BE and club cells (Club cell cluster 0 and BE cluster 6) could be distinguished from other cell states in the differentiation trajectory. Given that BE cells showed upregulated stem cell signature scores (**Supplemental Figure 1a**), we used BE cells as the starting point and plotted the diffusion pseudotime trajectory on the partition-based graph abstraction (PAGA) initialized embedding with a list of cell type specific markers as well as proliferation markers *MKI67* and *TOP2A* (**Supplemental Figure 4a,b**). We observed that *KRT5*+ BE cells gave rise to all other epithelial cells and tumor cells, with tumor cells and LE cells (*KLK3*+) appearing later than club cells (*PIGR*+, *LTF+* and *PSCA+*) in the pseudotime trajectory (**Supplemental Figure 4c**), consistent with a previous analysis^9^. We ran Monocle3^45^ to compute the pseudotime trajectory for PCa club cells (**Supplemental Figure 4d,e**). Club cells with higher LE signature scores were more differentiated in pseudotime (**Supplemental Figure 4e**). This finding was further supported by increasing expression levels of LE markers *ACPP* and *KLK3* along the trajectory compared to club cell markers (**Supplemental Figure 4f,g)**, suggesting that LE-like club cells in PCa samples could be transitioning to LE cells or tumor cells.

### Integrated epithelial cell analysis reveals upregulated *AR* signaling in PCa samples

As PCa samples in this study included four paired tumor and normal samples, we tested if PCa-enriched cell states in BE, LE, and club cells were enriched in the surrounding epithelial cells of the PCa biopsies and in radical proctectomy tissue samples containing tumor cells. We compared the percentage composition of each BE and club cell state within all BE and club cells in all five sample types respectively (Normal, biopsy, RP paired tumor, RP paired normal, and RP unpaired tumor). The PCa-enriched cell states of Club cell cluster 0 and BE cluster 6 were similarly represented in the four paired tumor and normal samples (p = 0.43, Mann-Whitney U test).

To identify the overall epithelial cell transcriptional programs in PCa samples, we integrated all PCa epithelial cells (Epithelial PCa) with prostate epithelial cells from normal healthy controls (Epithelial Normal)^9^ (**Figure 5a**). We identified differentially expressed genes between tumor and normal samples across all three major types of epithelial cells (LE, BE, and club cells). We found ATF transcription factors *FOS* and *JUN*, members of the EGFR pathway that mediate gene regulation in response to cytokines and growth factors^46^, and prostate acid phosphatase (*PSAP*)^47^ as commonly upregulated across these cell types (**Figure 5b**). However, the DEGs could not be recapitulated when comparing between paired tumor and normal samples (**Supplemental Table 5**), suggesting that compared to normal prostate samples, epithelial cells in the paired normal tissues were more similar to those from paired tumor tissues taken from different anatomical regions within the same radical prostatectomy specimen. Since the two PCa-enriched cell states in BE and club cells showed upregulated *AR* signaling compared to other BE or club cells respectively, we tested *AR* expression in the integrated dataset and found that in PCa epithelial cells, 21.4% of BE cells (458 of 2,145 cells), 28.6% of club cells (249 of 872), 52.7% of LE cells (2,974 of 5,647 cells) and 43.2% of tumor cells (1,993 of 4,658 cells) were *AR*+, in which significantly higher percentages of PCa BE, LE, and club cells were *AR*+ compared to the same cell types from normal samples (p < 0.001, FET) (**Figure 5c**). We also computed the Hallmark Androgen Response pathway signature scores for all cells and found that the three major epithelial cell types in PCa samples were all upregulated in *AR* signaling compared to normal samples (p < 0.001, Wilcoxcon rank sum test) (**Figure 5c**).

**Figure 5.**
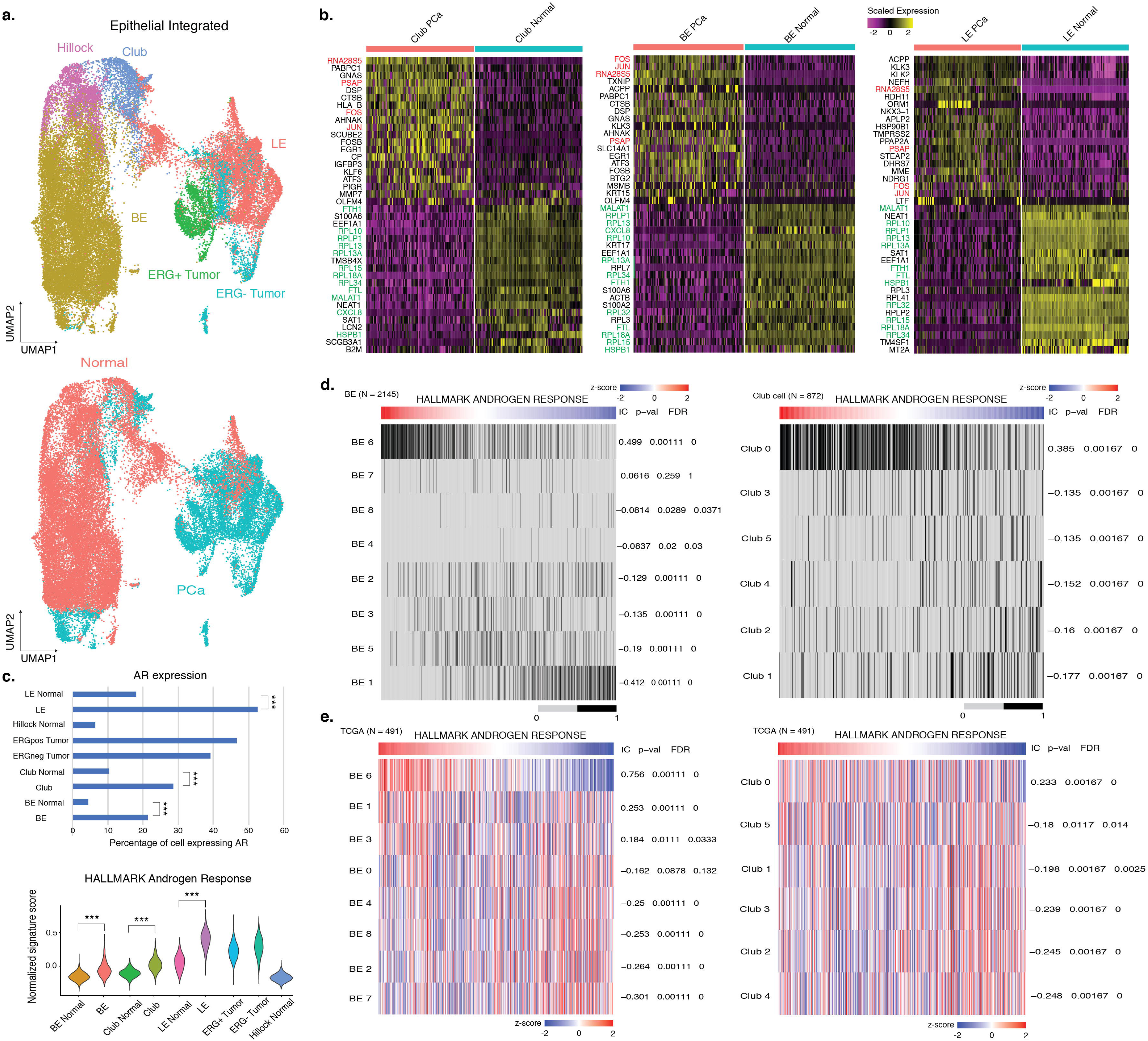
Integration of PCa and normal epithelial cells reveals common *AR* signaling upregulation driven by PCa-enriched BE and club cell states. **a.** UMAP of integrated epithelial cells annotated by cell types and sample type (PCa and Normal), then separated by the origin (either previous normal epithelial cells or epithelial cells in the PCa samples). **b.** Heatmaps of top 20 differentially expressed genes between PCa samples and normal prostates for adjacent cell types (left: BE PCa, BE Normal. Middle: Club Normal, Club PCa. Right: LE PCa, LE Normal). Commonly upregulated genes in the PCa samples are labeled in red, and commonly upregulated genes in the normal samples are labeled in green. **c.** Top, *AR* expression percentages in all epithelial cell types within the integrated dataset. Significance levels are labeled in each comparison (***: p < 0.001, FDR). Bottom, Comparison of Hallmark *AR* pathway signature scores of each epithelial cell type. Significance levels are labeled for each common cell type (*: p < 0.05, ***: p < 0.001, Wilcoxon rank sum test). **d.** The association of *AR* signature with BE and club cell state. Each cell is labeled (grey: 0, not in the cell state; black: 1, in the cell state). Information coefficient, accompanied p-values and FDR q values are labeled next to each cell state. **e.** The association of *AR* signature with BE and club cell state signature scores in the TCGA datasets (N = 491). Information coefficient, accompanied p-values and FDR q values are labeled next to each cell state.

To validate the two PCa-enriched epithelial cell states we identified in BE and club cells and test their correlation with upregulated *AR* signaling, we ran ssGSEA on all BE and club cells on the Hallmark Androgen Response pathway. The *AR* signature score of BE cells was only significantly positively correlated to BE cluster 6 (information coefficient (IC) = 0.499, FDR q < 1e-5), and the *AR* signature score in club cells was significantly positively correlated to Club cell cluster 0 (IC = 0.385, FDR q < 1e-5) (**Figure 5d**). Furthermore, to test if this correlation between a PCa-enriched cell state and *AR* signaling could be replicated in other PCa datasets, we projected all BE and club cell states across the TCGA^25^ (N = 499) and SU2C^48^ (N = 266) castration resistant prostate cancer (CRPC) bulk RNA-seq datasets (**methods**). In both bulk RNA-seq datasets, *AR* signature scores were positively correlated with BE cluster 6 (IC = 0.756, FDR q < 1e-5) and Club cell cluster 0 (IC = 0.233, FDR q < 1e-5) (**Figure 5e**), supporting our identification of cell states within BE cells and club cells that were more androgen responsive and associated with prostate cancer.

### Transcriptomic profiles of *ERG*+ tumor cells are patient-specific while *ERG*- tumor cells overlap with surrounding LE cells

While *ERG+* tumor cells clustered separately from non-malignant LE cells, *ERG-* tumor cells resided more closely to non-malignant LE cells (**Figure 2c**). To investigate this further, we first analyzed the sub-structure of *ERG+* and *ERG-* tumor cells separately to identify distinct underlying cell states (**Figure 6a,b**). *ERG+* tumor cells clustered in a patient-specific manner, whereas no such pattern was seen for *ERG-* tumor cells as most *ERG*- tumor cell states were comprised of more than one patient (**Figure 6c**).

**Figure 6.**
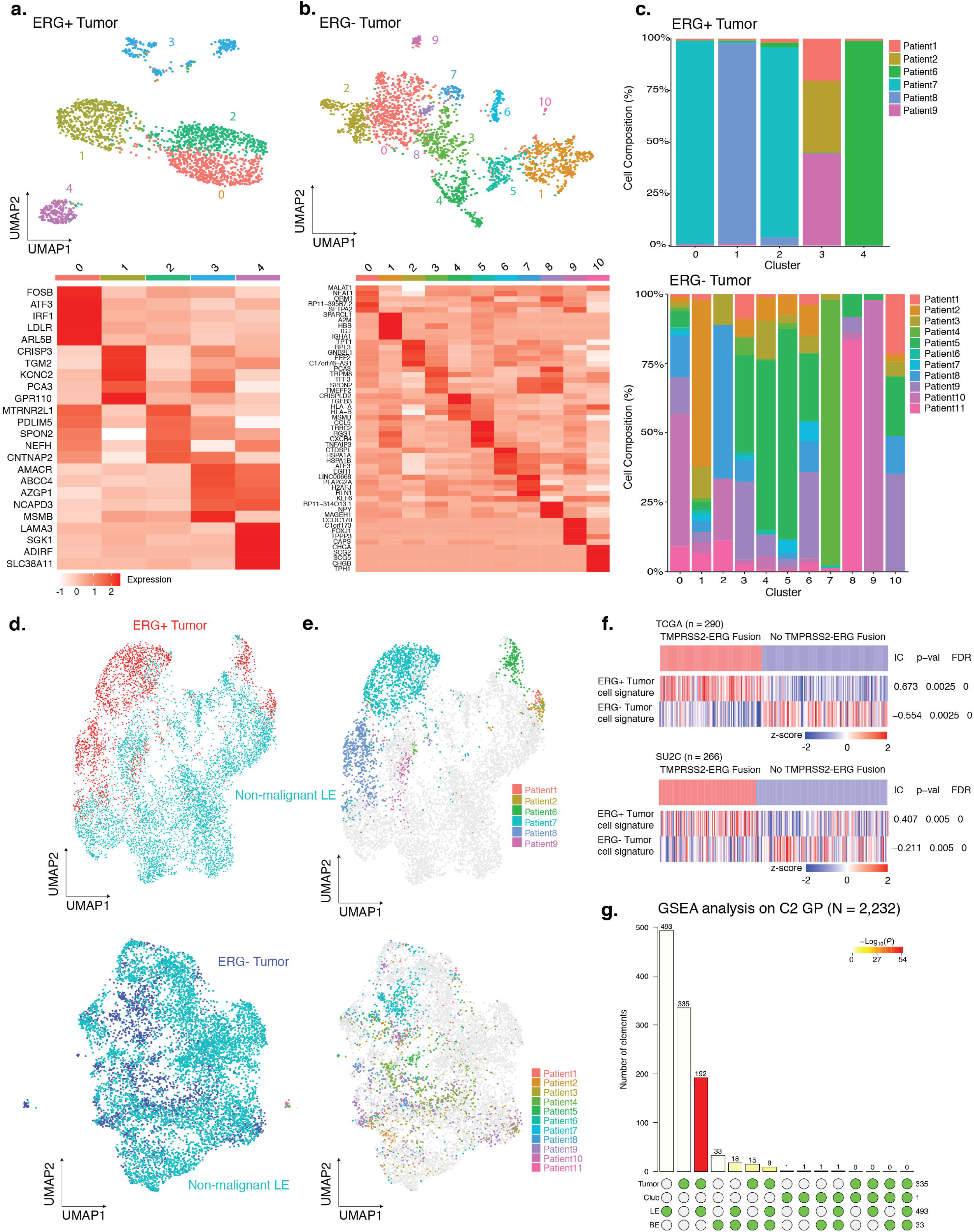
Comparison of *ERG*+ and *ERG*- tumor cells reveals patient-specific cell states and intra-patient heterogeneity. **a.** UMAP of *ERG*+ tumor cells labeled by clusters with differential gene expression profiles (top). Heatmap of the top 10 differentially expressed genes for each cluster (bottom). **b.** UMAP of *ERG*- tumor cells labeled by clusters with differential gene expression profiles (top). Heatmap of the top 10 differentially expressed genes for each cluster (bottom). **c.** Patient composition in each cluster for *ERG*+ tumor cells (top) and *ERG*- tumor cells (bottom). Cell counts in each cluster are normalized to 100%. **d.** UMAP of *ERG*+ and *ERG*- tumor cells when integrated with non-malignant LE cells respectively. **e.** UMAP of *ERG*+ and *ERG*- tumor cells when integrated with non-malignant LE cells labeled by patients. **f.** The association of *TMPRSS2-ERG* fusion status in the TCGA (N = 290) and SU2C (N = 266) datasets with *ERG*+ and *ERG*- tumor cell signature (red: *TMPRSS2-ERG* fusion detected; blue: *TMPRSS2-ERG* fusion not detected). Information coefficient, accompanied p-values and FDR q values are labeled. **g.** Visualization of the intersection amongst significant GSEA results for BE, LE and club cells. The color intensity of the bars represents the p- value significance of the intersections.

One possibility for the different distribution patterns between *ERG*+ and *ERG*- tumor cells is that *ERG*+ tumor cells for each patient represented a distinctive cell state driven by a dominant oncogenic alteration, though no such distinction was seen in *ERG*- tumor cells, suggesting more overlapping cell states between *ERG*- tumor cells and adjacent non-malignant LE cells. To test this hypothesis, we integrated *ERG*+ tumor cells and *ERG*- tumor cells separately with LE cells and performed sub-clustering analyses. Overall, we found 1,244 genes significantly varied between *ERG*+ tumor cells and LE cells (FDR q < 0.01, Wilcoxon rank sum test), while only 314 genes were significantly varied between *ERG*- tumor cells and LE cells (FDR q < 0.01, Wilcoxon rank sum test). Fourteen and seventeen cell states were recovered in the *ERG*+ and *ERG*- integrated datasets, respectively (**Supplemental Figure 5a-b**). We observed a clear separation between *ERG*+ tumor cells and non-malignant LE cells while *ERG*- tumor cells were not clearly distinguishable from non-malignant LE cells in the analysis (**Figure 6d**). From the cell state composition comparison, we observed three cell states with more than 400 cells each that were almost exclusively detected in the *ERG*+ tumor cells, with each cell state largely attributed to one specific patient (**Supplemental Figure 5a**). In contrast, no such patient specificity was observed for *ERG-* tumor cells (**Figure 6e**) (**Supplemental Figure 5b**). In our dataset, *ERG*+ tumor cells were predominantly found in tumor samples while *ERG*- tumor cells were found in paired tumor and normal samples (**Supplemental Figure 5c**). Using the DEGs between *ERG*+ and *ERG*- tumor cells (**Supplemental Figure 5d**), we generated signature gene sets for both types of tumor cells and tested if the signatures of *ERG*+ and *ERG*- tumor cells generated from this dataset were correlated with *TMPRSS2-ERG* fusion status in TCGA^25^ and SU2C^48^ castration resistant prostate cancer (CRPC) bulk RNA-seq datasets. *TMPRSS2-ERG* fusion status was significantly positively correlated with an *ERG*+ tumor cell signature score in both datasets (TCGA: information coefficient (IC) = 0.673, FDR q < 1e-5; SU2C: IC = 0.407, FDR q < 1e-5) and the absence of *TMPRSS2- ERG* fusion was significantly correlated with *ERG*- tumor signature scores (TCGA: IC = -0.554, FDR q < 1e-5; SU2C: IC = -0.211, FDR q < 1e-5) (**Figure 6f**). These results supported the tumor cell signatures and our use of *ERG* expression as a classification in annotating tumor cells.

Furthermore, we compared the numbers of *ERG*+ tumor cell and *ERG-* tumor cells in each patient. Tumor cells in five patients were over 90% *ERG*- and over 90% *ERG*+ in two patients (**Supplemental Figure 5e**) Tumor cells in four patients harbored both types of tumor cells. Using non-tumor epithelial cells as reference, we found significantly different CNV profiles from the reference for each patient, further validating our tumor cell identification (**Supplemental Figure 5f**). For our downstream analyses, we classified patients based on ERG status by annotating the five patients with almost exclusive *ERG*- tumor cells as *ERG*- patients and the other six patients (exclusive *ERG*+ tumor cells and mixtures) as *ERG*+ patients.

### T-cell and stromal cell analysis reveals common signaling in *ERG*- patients

The transcriptional differences between *ERG*+ and *ERG*- tumor cells suggested that they might give rise to differential responses in the tumor microenvironment. To identify tumor-related immune cells and whether specific immune cell types were differentially enriched in either *ERG*+ or *ERG*- samples, we analyzed the T-cell population and identified CD4 and CD8 T-cells, regulatory T-cells (Treg), and NK cells based on differentially expressed genes (**Figure 7a**). We then stratified the T-cell populations based on *ERG* status and found two CD4 T-cell clusters that were differentially enriched. Between the two CD4+ T-cells we identified, CD4 T-cell cluster 1 was enriched in *ERG*+ patients with a 2.73 fold difference (20.5% vs 7.5%) (**Figure 7b**) and was characterized by a higher level expression of immune response regulators including AP-1 transcriptor factors^49^ *FOSB* (log2FC = 1.79, FDR q = 5e-30), *FOS* (log2FC = 1.78, FDR q = 6.2e-26) and *JUN* (log2FC = 1.55, FDR q = 5.5e-22). CD4 T-cell cluster 2 was enriched in *ERG*- patients with a 5.6 fold change (9.5% vs 1.7%) (**Figure 7b**) (p < 0.001, Fisher’s exact test) and was marked by higher expression of *DUSP4* (log2FC = 1.30, FDR q = 1.4e-20) and *CXCR6* (log2 fold change (log2FC) = 1.31, FDR q = 1.5e-22), which was previously shown to be expressed in the type-1 polarized T-cell subset and to contribute to tumor progression^50^. We noted that the DEGs between the two T-cell clusters were consistent with the DEGs identified between *ERG*+ and *ERG*- tumor cells, with *FOSB, FOS,* and *JUN* overexpressed in *ERG*+ tumor cells while *CXCR6* and *DUSP4* were overexpressed in *ERG*- tumor cells (**Supplemental Figure 5d**). No other T-cell populations (CD8 T-cells, Treg, and NK cells) showed a significant difference in cell type abundance between *ERG*+ and *ERG*- patients.

**Figure 7.**
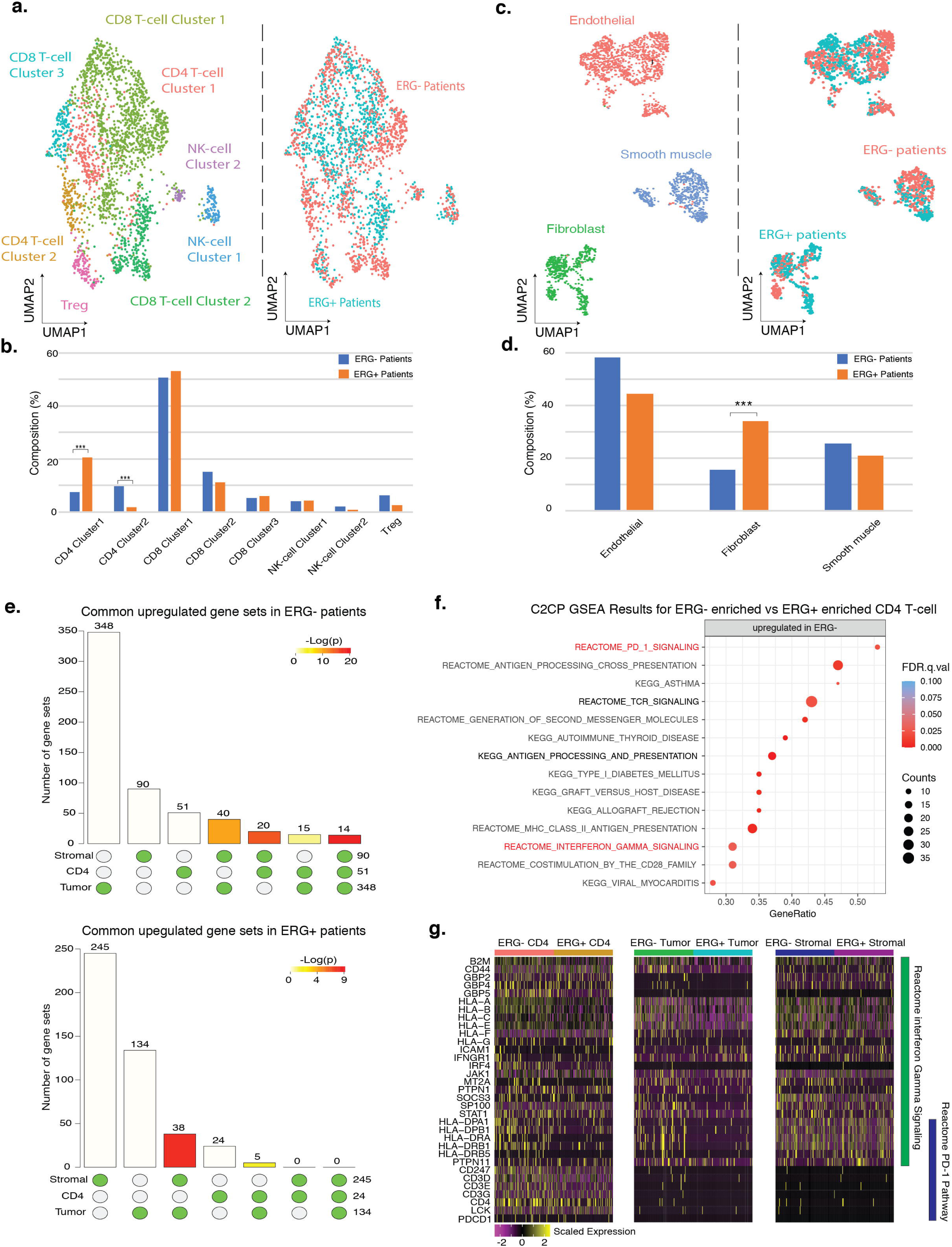
CD4 T subsets associated with *ERG* status and common upregulation of PD-1 and interferon gamma signaling in the *ERG*- tumor microenvironment. **a.** UMAP of T-cells labeled by different cell types (left) and *ERG*+ or *ERG*- patients (right). **b.** Cell composition comparison between *ERG*+ and *ERG*- patients for all T-cell cell types. Significance levels are labeled in differentially enriched clusters. **c.** UMAP of stromal cells labeled by different cell types (left) and *ERG*+ or *ERG*- patients (right). **d.** Cell composition comparison between *ERG*+ and *ERG*- patients for all stromal cell types. Significance levels are labeled in differentially enriched clusters. **e.** Visualization of the intersections amongst significantly upregulated (top) and downregulated (bottom) gene sets within C2 CP gene set collection for tumor cells, two clusters of differentially enriched CD4 T-cell clusters and stromal cells. Significant GSEA results are represented by circle below bar chart with individual blocks showing “presence” (green) or “absence” (grey) of the gene sets in each intersection. P-value significance of the intersections are represented by color intensity of the bars. **f.** GSEA results for the *ERG*- patient-enriched CD4 T-cell cluster compared to the *ERG*+ patient-enriched cluster on the common upregulated gene sets (N = 14). Gene counts for the corresponding gene set are indicated by marker radius. Statistical significance levels (FDR) are shown by color gradient. Reactome PD-1 and Interferon gamma signaling pathways are highlighted in red. **g.** Gene expression heatmaps of genes in the Reactome PD-1 and Interferon gamma signaling pathways for tumor cells, CD4 T-cells and stromal cells in both *ERG*+ and *ERG*- patients.

Similarly, we stratified the stromal population based on the *ERG* status of patients and identified three distinct clusters consistent with endothelial cells, smooth muscle cells, and fibroblasts (**Figure 7c**). Of these three stromal cell types, fibroblasts showed an enrichment in *ERG*+ patients (p < 0.001, FET).

To test if the differences between *ERG*- and *ERG*+ tumor cells could potentially drive distinct and common stromal and immune responses, we ran independent GSEA analyses between *ERG*- and *ERG*+ tumor cells, CD4 T-cells and stromal cells and computed the intersection of significantly upregulated gene sets in *ERG*- patients (FDR q < 0.1). Fourteen upregulated gene sets were identified that were commonly upregulated in *ERG*- tumor cells, CD4 T-cells and stromal cells (p < 10e-20, multi-set intersection exact test^51^) (**Figure 7f**). However, we did not detect any common pathway changes in the other epithelial populations (**Figure 6g**). The fourteen common upregulated gene sets in *ERG*- patients included Reactome PD-1 and Reactome interferon gamma signaling (**Figure 7g**), which have both been reported to be upregulated in advanced prostate cancers^52, 53^. Within these two gene sets, we found that *ERG*- patient-enriched CD4 T-cells, tumor cells, and stromal cells showed significantly higher expression of a family of HLA genes compared to *ERG*+ cell populations (p < 0.05, Wilcoxon rank sum test) (**Figure 7h**). Within the T-cells, while there was no difference in the cell composition of CD8 T-cells based on *ERG* status, the *ERG*- CD8 T-cell population was also found to be upregulated in the Reactome PD-1 and Reactome interferon gamma signaling signatures (FDR q < 0.1, **Supplemental Table 4**). To test if *ERG*- tumor cell-associated CD4 and CD8 T-cells could represent a distinct immune cell niche, we tested a series of exhausted, cytotoxic markers^54^ as well as genes in the PD-1 and Reactome interferon gamma signaling pathway (**Supplemental Table 6**). We found that *ERG*- CD4 T-cells were significantly upregulated in exhausted T-cell markers *PDCD1* (log2FC = 0.52, p < 0.01, Wilcoxon rank sum test) and *CTLA4* (log2FC = 1.79, p < 0.001, Wilcoxon rank sum test) and cytotoxic markers *GZMA* (log2FC = 1.54, p < 0.001, Wilcoxon rank sum test) and *GZMB* (log2FC = 1.09, p < 0.05, Wilcoxon rank sum test) compared to *ERG*+ CD4 T-cells. Additionally, *ERG*- CD8 T-cells were upregulated in exhausted T-cell markers *HAVCR2* (log2FC = 0.68, p < 0.05, Wilcoxon rank sum test) and *LAG3* (log2FC = 0.86, p < 0.001, Wilcoxon rank sum test) compared to *ERG*+ CD8 T-cells (**Supplemental Figure 6a,b**). These results suggested that CD4 and CD8 T-cells associated with *ERG*- tumor cells represented a more exhausted and cytotoxic phenotype. Then, using CD4 phenotype markers from a previous analysis^55^, we tested the frequency of expression for these markers in both *ERG*+ and *ERG*- CD4 T-cells and found a significantly higher proportion of *CCR7*+ central memory CD4 T-cells, *CD69*+ activated CD4 T-cells, *GZMB*+ cytotoxic CD4 T-cells, and *TOX*+ exhausted CD4 T-cells^55^ associated with *ERG*- patients (**Supplemental Figure 6c**).

After T-cells, myeloid cells comprised the second largest immune cell population. Annotation of the myeloid cell population with SingleR^19^ yielded four cell types: neutrophils, eosinophils, macrophages, and monocytes (**Supplemental Table 7**; **Supplemental Figure 7a-b**). Within the myeloid cell population, we did not detect any significant composition differences in monocytes or macrophages between RP paired tumor and paired normal samples or between *ERG*+ and *ERG*- patients (p > 0.05, FET) (**Supplemental Figure 7c**).

To investigate the subtypes of monocytes and macrophages that are associated with tumor-related responses, we identified monocytes and macrophages with high expression of cell cycle markers *MKI67* and *TOP2A*, indicating a cluster of proliferating myeloid cells (**Supplemental Figure 7d**) that we termed *MKI67*+ myeloid cells. Monocytes were further classified by the expression of *CD14* (**Supplemental Figure 7d)**. Within the macrophage population, we used previously established signatures^56–59^ of dichotomous phenotypes to classify macrophages into M0, M1, and M2 types, of which M1 macrophages have been described as pro-inflammatory and M2 macrophages as anti-inflammatory and associated with tumor progression^60^. We computed the signature scores of M1 and M2 macrophages and annotated the two subtypes accordingly, based on signature scores as well as M1 specific markers, such as *IL1A*, *CXCL3,* and *PTGS2*, and M2 specific markers, such as *ARG1*, *CCL22,* and *FLT1*. Neither M1 nor M2 macrophages were clustered separately from normal M0 macrophages, consistent with a previous analysis of macrophage subtypes^58^ (**Supplemental Figure 7d,e**).

A recent study on macrophages categorized macrophages into resident tissue macrophages enriched in normal tissues (RTM) and tumor associated macrophages (TAM) enriched in tumor tissues, which did not fit the M1/M2 phenotypes^61, 62^. We did not detect RTMs within the PCa samples (**Supplemental Figure 7f**). In contrast, TAMs were described as either *C1QC*+ or *SPP1*+. These TAMs were reported to derive from *FCN1*+ monocyte-like macrophages, which was consistent with the detection of *FCN1* in a cluster of PCa myeloid cells where we saw a mixture of monocytes and macrophages (**Supplemental Figure 7f**). In total, 713 TAMs were identified but no significant difference in composition was detected between paired tumor and normal samples (77.9% vs 69.0%, p = 0.58, FET) (**Supplemental Figure 7g**).

Another group of tumor-associated myeloid cells termed myeloid-derived suppressor cells (MDSC) has been characterized with roles in inflammation, establishing host immune homeostasis, and driving castration resistance in prostate cancer^63–66^. These MDSCs can inhibit anti-tumor reactivity of T-cells and NK-cells and the enrichment of MDSCs was correlated with tumor progression and worse clinical outcomes^67^. Two types of MDSCs have been described: monocytic MDSC (M-MDSC) characterized by high expression of *CD11* and *CD14* and low expression of *HLA* and *CD15* and granulocytic or polymorphonuclear MDSC (PMN-MDSC) characterized by high expression of *CD11* and *CD15* and low expression of *CD14*. To test for the presence of these MDSCs in our PCa samples, we used the co-expression of these markers and identified 137 M-MDSCs within the 790 *CD14+* monocytes and 11 PMN-MDSCs within 974 *CD14-* monocytes (**Supplemental Figure 7g)**. M-MDSCs were enriched in the paired tumor samples compared to paired normal (19.9% vs 3.6% of total monocytes, p = 0.0035, FET).

### Prostate cancer organoids recapitulate epithelial cell types with uniquely expanded cell states in BE and club cells

To develop models to examine the cellular state heterogeneity revealed by single-cell analysis and to determine if we could reconstitute and propagate prostate cancer-associated club cells, we used established methods^68, 69^ to generate localized prostate cancer organoids from single cells from six patients who underwent radical prostatectomies (four patients included in the tissue sample dataset) and characterized them using scRNA-seq within three passages (**Figure 8a**). PCA-based clustering of organoid samples yielded 23 clusters from a total of 15,073 cells. We identified a total of six epithelial cell types with distinctive DEGs, based on the cell type signatures we generated from the PCa tissue samples and the established signatures from normal samples **(Supplemental Table 2)** (**Figure 8a**). The epithelial cell types included BE cells characterized by high expression of *DST*, *KRT15, KRT5, KRT17,* and *TP63*, club cells characterized *by PIGR, MMP7, CP,* and *CEACAM6,* hillock cells, consistent with those in normal prostates showing high level expression of *KRT13, CLCA4,* and *SERPINB3*, a mesenchymal stem cell (MSC) population expressing known MSC markers^70–72^ *LAMC2*, *VIM, MMP1,* and *KLK7* and a population with high level expression of cell cycle markers *MKI67* and *TOP2A* termed *MKI67*+ epithelial cells (**Supplemental Figure 8a**). Notably, within these early passage organoids we identified a tumor cell population expressing a high level of LE cell markers (*KLK3, KLK2,* and *ACPP*) and tumor markers (*PCA3, TRPM8,* and *ERG*) (**Supplemental Figure 8a**). Cell type annotation was supported by ssGSEA, which showed that the MSC population was upregulated in the MSC signature gene set developed from a previous analysis^71^ and that the *MKI67+* cluster was upregulated in a KEGG cell cycle signature indicating proliferating cells (**Supplemental Figure 8b**). The identification of tumor cells was further validated by InferCNV^21^ estimation (**Supplemental Figure 8c**). To validate our recovery of the cell type diversity in the organoids, we performed immunofluorescence staining for *KRT8*+ luminal and *KRT5*+ basal cells (**Figure 8b**). We validated club cell proliferation *in vitro* by staining for *SCGB1A1*, an established club cell marker in the lung and prostate^9^, and lactoferrin (*LTF*), which was upregulated in the PCa club cells identified by scRNA-seq (**Figure 8b**).

**Figure 8.**
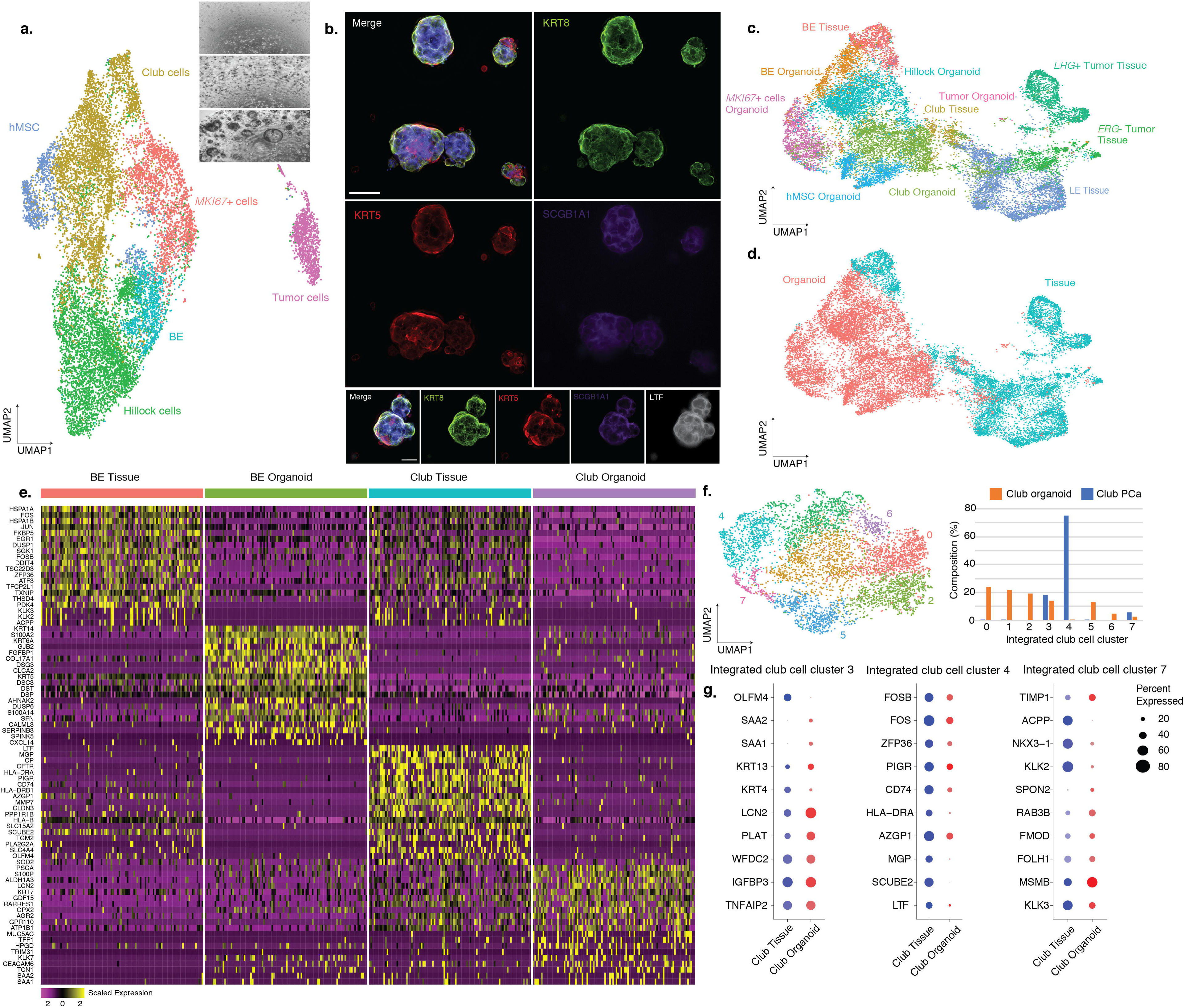
*In vitro* organoid samples recapitulate PCa-enriched BE and club cell states. **a.** UMAP of cells from organoid samples labeled by different cell types. Organoid culture snapshots are depicted in the upper right panel. **b.** Immunofluorescence staining for LE marker (*KRT8*), BE marker (*KRT5*) and club cell markers (*SCGB1A1, LTF*) of the organoid samples. **c.** UMAP of integrated dataset of cells from the organoid samples and epithelial cells from matching parent tissue samples, labeled by cell types. **d.** UMAP of integrated dataset, labeled by sample types (tissue or organoid samples). **e.** Heatmaps for the top 20 differentially expressed genes for BE and club cells between tumor tissues and organoid samples. **f.** UMAP of integrated club cell dataset of tumor tissue and organoid samples. Cell composition comparison is shown in the grouped bar charts. **g.** Dot plots of the top 10 differentially expressed genes in cluster 3, 4 and 7 in tissue and organoid club cells. Dot size represents proportions of gene expression in cells and expression levels are shown by color shading (low to high reflected as light to dark).

To test the fidelity of the organoids as models for tumor tissues, we integrated the cells in the early-passage (P0-P3) organoid samples (N = 10,990) with the epithelial cells from the four RP specimens from which the organoids were derived (N = 8,719) (**Figure 8c**). Compared to PCa tissue samples, LE cell markers or signature scores could not identify a distinctive LE cell cluster in the organoid samples (**Supplemental Figure 8b**), consistent with a previous study that LE cells were rarely captured in *in vitro* organoid cultures analyzed by scRNA-seq^73^. For the four patient-derived organoids, only a small number of tumor cells were captured compared to the parent tissues (tissue samples vs organoids, 34.11% vs 0.11%). However, hillock cells, MSCs and a population of *MKI67*+ epithelial cells were exclusive to the organoid samples and were not observed in PCa tissue samples (**Figure 8d**).

As BE and club cells were the two primary overlapping cell types between tissue and organoid samples (representing 11.9% and 29.0% of all cells, respectively, in the organoid samples), we took the subset of BE cells and club cells in tissue and organoid samples from the integrated dataset and computed the DEGs. BE markers *KRT5*, *DST,* and *KRT15* were expressed in BE populations from tissue and organoids and club cell genes *MMP7*, *LCN2,* and *CP* were expressed in both club cell populations (**Figure 8e**), suggesting similarities between tissue and organoid BE and club cells.

We then investigated BE and club cell populations by integrating organoids with tissue samples, respectively, to identify cell state differences in the organoid samples. We identified nine clusters in the integrated BE cell dataset with distinctive groups of DEGs (**Supplemental Figure 8d**). Compared to BE cells in PCa tissue samples, significantly higher percentages of BE cells in organoids expressed *KRT6A* (organoid vs tissue, 77.4% vs 0.56%, p < 0.001, FET)*, KRT14* (organoid vs tissue, 71.2% vs 18.6%, p < 0.001, FET), and *KRT23* (organoid vs tissue, 78.8% vs 20.2%, p < 0.001, FET) (**Supplemental Figure 8e**), suggesting that BE cells in organoid samples may be more representative of a progenitor cell state.

Similarly, when analyzing the organoid club cells with club cells from PCa tissues, we identified a total of eight clusters with distinctive DEGs (**Supplemental Figure 8f**) and observed an expansion of cell states in the organoid samples (**Figure 8f**). Among the eight clusters, five were predominantly comprised of organoid club cells, while club cells from prostate tissue were only found in clusters 3, 4 and 7. By comparing the expression levels of the top differentially expressed genes for these three clusters split by tissue and organoid club cells, we found that in cluster 3, hillock cell marker *KRT13* was expressed in tissue and organoid club cells, suggesting an intermediate hillock-club cell state. In cluster 4, PCa club cell marker *PIGR* was detected in 47% of organoid club cells (16 of 34) and 71% of tissue club cells (325 of 653). *LTF* was expressed in 15% of organoid club cells (5 of 34) compared to 50% tissue club cells (326 of 653), suggesting that *LTF* may be a PCa tissue-specific club cell marker. In contrast, the top DEGs for cluster 7 included LE markers such as *ACPP, NKX3-1, KLK2* and *KLK3*, consistent with the profile of the previously-identified PCa-enriched club cell state (**Figure 8g**). In cluster 7, we observed approximately 20% of organoid club cells expressing at least one LE cell marker. This cluster scored higher for the PCa-enriched club cell state compared to all other clusters of organoid club cells, suggesting that PCa-enriched club cell states were recapitulated in organoid samples.

Overall, we found that organoid samples harbored cell states found in tumor tissues and an enrichment of progenitor-like cell states and intermediate cell states. The plasticity of these organoid-enriched cell states within BE and club cells suggests that *in vitro* organoid models may provide useful models to study cell state differences and identify lineage relationships to tumorigenesis.

## Discussion

Studies of localized prostate cancer have been extensively performed with bulk RNA-seq and WES/WGS approaches that have provided key insights into the molecular features of prostate cancer^9, 12, 63–66^. Here, we performed single-cell analyses of localized PCa biopsies and radical prostatectomy specimens to characterize the heterogeneity of tumor cells and subpopulations of epithelial cells, stromal cells, and tumor microenvironments.

Of note, we identified a distinctive epithelial cell population of club cells that has not been previously observed in human prostate cancer samples. While club cells have been noted in normal prostates^9, 77, 78^, a population of club cells associated with prostate cancer suggests they may play a previously unappreciated role in carcinogenesis. Recent studies have identified a progenitor-like *CD38*^low^*/PIGR*^high^*/PSCA*^high^ luminal epithelial cell sub-population with regenerative potential^39, 78, 79^. Based on the similarity of highly expressed genes including *PIGR*, *MMP7*, *CP,* and *LTF*, we believe those cells are consistent with their identity as club cells. In our analysis, prostate cancer club cells are characterized by the markedly lower expression of *SCGB3A1* and *LCN2* compared to club cells from normal healthy controls^9^. Based on our gene signature analyses, our results suggest that PCa club cells are more androgen responsive overall and harbor a highly androgen-responsive cell state that may be a potential progenitor cancer cell or function to support the overall androgen responsive cellular milieu of prostate cancer^80, 81^.

*SCGB3A1*, a marker for club cells, was one of the top downregulated genes in prostate cancer club cells compared to club cells from normal healthy control prostates. *SCGB3A1* may play a tumor suppressor role in a number of cancers including breast, prostate, and lung as its expression has been noted to be markedly lower in cancer tissues compared to normal tissues^82^. We speculate that prostate club cells in the normal epithelia may play a tumor suppressor role through secretion of *SCGB3A1* which is then downregulated in concert with prostate cancer progression, as marked by our finding of *SCGB3A1*^low^ club cells in prostate cancer tissues that can be propagated in organoids. We did not find a distinct population of hillock cells in prostate cancer tissues so it is possible that hillock cells may be depleted in prostate cancer progression.

Consistent with other cancer single cell studies in which tumor cells cluster separately, *ERG*+ tumor cells clustered separately by patient from non-malignant epithelial clusters^14, 54, 83–85^. However, our analysis of *ERG*- tumor cells unexpectedly found that *ERG*- tumors did not cluster by patient and we observed a shared heterogeneity for *ERG*- tumor cells with non-malignant luminal cells.

Treating prostate cancer with immune checkpoint inhibition has had limited efficacy to date and these therapies have largely focused on advanced castration-resistant tumors^43, 44, 86–90^. Our single-cell analysis reveals new insights into the tumor immune microenvironment of localized prostate cancer based on *ERG* status. We hypothesized that *ERG*- tumor cells might evoke similar tumor microenvironment responses and found common transcriptional pathways that were upregulated in the tumor, stroma, and CD4 T cell populations of *ERG*- patients, including the PD-1 and interferon gamma signaling pathway, suggesting that *ERG*- tumor cells may give rise to a distinct immune cell niche and tumor microenvironment.

We note a potential limitation of our analysis in the identification of *ERG*- tumor cells as we also found evidence for *ERG*- tumor cells in paired grossly normal specimens. This could be attributed to tumor cells also being present in the seemingly normal tissues from radical prostatectomy specimens^14, 85, 91, 92^. Analysis of somatic mutations or structural variants on a single-cell level will contribute to the identification of *ERG-* tumor cells and inform our understanding of tumor heterogeneity.

Furthermore, we showed that *in vitro* organoid cultures grown from tumor specimens can recapitulate cell states found in tumor tissues. We identified a number of new cell types that emerged in the organoid samples including hillock cells, MSC and *MKI67*+ epithelial cells. The mechanisms by which hillock cells can propagate in organoid cultures but not be found in the localized tumor tissue specimens are still to be delineated. An expansion of cell states in BE and club cells in the organoids suggests a broader view for their capacity for cell state transitions. Our results suggest that prostate cancer epithelial organoids harbor many major cell types from tissue and provide a useful model to investigate cell state plasticity in the context of selective pressures and genetic perturbations. However, in contrast to previous studies on organoids generated from prostate samples, we did not observe a distinctive *NKX3-1*+/*KLK3+/AR+* luminal cell population^68, 93, 94^. This might be due to a limitation of detection using single cell sequencing technology or that we could not robustly grow differentiated luminal cells^73^.

Comparing epithelial cells from PCa samples with those from normal healthy controls revealed distinct high androgen-signaling cell states that were enriched in PCa samples. We found that epithelial cells from PCa tissues were generally upregulated in *AR* signaling. Given our identification of shared luminal-like, highly androgen-responsive cell states across basal and club cell populations, we posit that these cell types may be primed for tumor cell transformation and may also promote prostate tumorigenesis. Further studies with lineage tracing and dissection of single cell somatic alterations within these specific cell states will be informative for further characterization of their potential tumorigenic roles. The identification of a tumor-associated club cell population raises the possibility that these cells contribute to the interactions between tumor cells and their surrounding epithelial microenvironment. Furthermore, our analyses identify cell type specific signature gene sets within prostate cancer samples that should contribute to a more precise and thorough classification of cells during prostate carcinogenesis. In summary, we provide a single-cell transcriptomic blueprint of localized prostate cancer that identifies and highlights the multicellular milieu and cellular states associated with prostate tumorigenesis. Our results provide new insights into the epithelial microenvironment and the cellular state changes associated with prostate cancer toward improved PCa diagnosis.

## Methods

### Experimental Details

#### Samples selection

We obtained a total of six prostate biopsies from three different patients (two biopsies for patient 1-3, obtained at the same time point), four radical prostatectomies with tumor-only samples from four patients (patients 4-7) and four radical prostatectomies with matched normal samples from four patients (patients 8-11, matched normal samples were taken from adjacent seemingly normal regions). Clinical/pathological data available for the samples is in **Supplemental Table 1.**

#### Study Approval

The UCSF Institutional Review Board (IRB) committee approved the collection of the patient data included in this study.

#### Tissue Dissociation

Tissue samples were minced with surgical scissors and washed with RP-10 (RPMI + 10% FBS). Each sample was centrifuged at 1200 rpm for five minutes, resuspended in 10 mL digestive media (HBSS + 1% HEPES) with Liberase TM (Roche, Cat: 5401119001) or 1000 U/mL collagenase type IV (Worthington, Cat: LS004188), and rotated for 30 minutes at 37 °C. Samples were triturated by pipetting ten times after every ten minutes during the incubation or by pipetting 15 times at the end of the incubation. Each sample was filtered through a 70 µm filter (Falcon, Cat: 352350), washed with RP-10, centrifuged at 1200 rpm for five minutes, washed again with RP-10, and resuspended in RP-10. A hemocytometer was used to count the cells.

#### Single-cell RNA sequencing

Sequencing was largely based on the Seq-Well S^3 protocol^13, 95^. One to four arrays were used per sample. Each array was loaded as previously described with approximately 110,000 barcoded mRNA capture beads (ChemGenes, Cat: MACOSKO-2011-10(V+)) and with 10,000-20,000 cells. Arrays were sealed with functionalized polycarbonate membranes (Sterlitech, Cat: PCT00162X22100) and were incubated at 37°C for 40 minutes.

After sealing, each array was incubated in lysis buffer (5 M Guanidine Thiocyanate, 1 mM EDTA, 0.5% Sarkosyl, 1% BME). After detachment and removal of the top slides, arrays were rotated at 50 rpm for 20 minutes. Each array was washed with hybridization buffer (2 M NaCl, 4% PEG8000) and then rocked in hybridization buffer for 40 minutes. Beads from different arrays were collected separately. Each array was washed ten times with wash buffer (2 M NaCl, 3 mM MgCl_2_, 20 mM Tris-HCl pH 8.0, 4% PEG8000) and scraped ten times with a glass slide to collect beads into a conical tube.

For each array, beads were washed with Maxima RT buffer (ThermoFisher, Cat: EP0753) and resuspended in reverse transcription mastermix with Maxima RT buffer, PEG8000, Template Switch Oligo, dNTPs (NEB, Cat: N0447L), RNase inhibitor (Life Technologies, Cat: AM2696), and Maxima H Minus Reverse Transcriptase (ThermoFisher, Cat: EP0753) in water. Samples were rotated end-to-end, first at room temperature for 15 minutes and then at 52°C overnight. Beads were washed once with TE-SDS and twice with TE-TW. They were treated with exonuclease I (NEB), rotating for 50 minutes at 37°C. Beads were washed once with TE-SDS and twice with TE-TW, and once with 10 mM Tris-HCl pH 8.0. They were resuspended in 0.1 M NaOH and rotated for five minutes at room temperature. They were subsequently washed with TE-TW and TE. They were taken through second strand synthesis with Maxima RT buffer, PEG8000, dNTPs, dN-SMRT oligo, and Klenow Exo- (NEB, Cat: M0212L) in water. After rotating at 37°C for one hour, beads were washed twice with TE-TW, once with TE, and once with water.

KAPA HiFi Hotstart Readymix PCR Kit (Kapa Biosystems, Cat: KK2602) and SMART PCR Primer were used in whole transcriptome amplification (WTA). For each array, beads were distributed among 24 PCR reactions. Following WTA, three pools of eight reactions were made and were then purified using SPRI beads (Beckman Coulter), first at 0.6x and then at a 0.8x volumetric ratio.

For each sample, one pool was run on an HSD5000 tape (Agilent, Cat: 5067-5592). The concentration of DNA for each of the three pools was measured via the Qubit dsDNA HS Assay kit (ThermoFisher, Cat: Q33230). Libraries were prepared for each pool, using 800-1000 pg of DNA and the Nextera XT DNA Library Preparation Kit. They were dual-indexed with N700 and N500 oligonucleotides.

Library products were purified using SPRI beads, first at 0.6x and then at a 1x volumetric ratio. Libraries were then run on an HSD1000 tape (Agilent, Cat: 50675584) to determine the concentration between 100-1000 bp. For each library, 3 nM dilutions were prepared. These dilutions were pooled for sequencing on a NovaSeq S4 flow cell.

The sequenced data were preprocessed and aligned using the dropseq_workflow on Terra (app.terra.bio). A digital gene expression matrix was generated for each sample, parsed and analyzed following a customized pipeline. Additional details are provided below.

#### Organoid culture

Isolated single cells not used for single-cell sequencing were additionally frozen in FBS + 10% DMSO, flash frozen on dry ice, or plated in Matrigel to grow as 3D prostate organoid cultures. Organoid cultures were established by plating 20,000 cells in 25uL Matrigel (Corning, Cat: 356231) in 48-well flat-bottom plates (Corning, Cat: EK-47102). Prostate-specific serum-free culture media contained 500 ng/mL human recombinant R-spondin (R&D Systems, Cat: 10820-904), 10uM SB202190 (Sigma, Cat: S7076), 1uM Prostaglandin E3 (Tocris, CAt: 229610), 1nM FGF10 (Peprotech, Cat: 100-26), 5 ng/mL FGF2 (Peprotech, CAt: 100-18B), 10 ng/mL 5alpha-Dihydrotestosterone (Sigma, Cat: D-073-1ML), 100 ng/mL human Noggin (Peprotech, Cat: 102-10C), 500nM A83-01 (Fischer, Cat: 29-391-0), 5 ng/mL human EGF (Peprotech, Cat: AF-100-15), 1.25mM N-acetyl-cysteine (Sigma, Cat: A9165), 10mM Nicotinamide (Sigma, Cat: N3376), 1X B-27 (Gibco, Cat: 17504044), 1X P/S (Gibco, Cat: 15140122), 10mM HEPES (Gibco, CAt: 15630080), and 2mM GlutaMAX (Gibco, Cat: 35050061)^69^. Additionally, 10uM Y-27 (Biogems, Cat: 1293823) was included during the first 2 weeks of growth and after passaging to promote growth^69^. Generally, organoid growth was apparent within two to three days and robust after two weeks. 250uL media was refreshed every two to four days using media stored at 4°C for a maximum of ten days. Organoid growth was monitored using an EVOS-FL microscope.

To passage prostate organoid cultures every 7-14 days, culture media was replaced with 300 uL TrypLE (1X, Gibco, Cat: 12604013). Individual domes were collected into 15mL Falcon tubes, disrupted by pipetting with wide-orifice tips and incubated at 37°C for 30 minutes. Following incubation, the dissociation media was neutralized using 10mL wash media: adDMEM/F12 containing 5% FBS, P/S, 10mM HEPES (1M, Gibco, Cat: 15630080) and 2mM GlutaMAX (100x, Gibco, Cat: 35050061)^69^. Cells were spun down at 500 G for five minutes and resuspended in 2mL wash media. Finally, the media was aspirated, cells were resuspended in Matrigel, and 25 uL/dome were plated per well.

Organoids were accessed using single-cell sequencing at an early passage (P0-4). To isolate single cells from Matrigel, organoids were collected in 500uL Trypsin (0.25%, Gibco, Cat: 25-200-056) and incubated at 37°C for 30-45 minutes until few clumps were visible. Throughout incubation, cells were triturated every five minutes. Single cells were resuspended in 9mL DMEM + 5% FBS + 0.05mM EDTA and passed through a 40 µM filter, followed by an additional wash of the filter with 1mL DMEM + 5% FBS + 0.05mM EDTA. Cells were spun down at 300 G for 5 minutes, resuspended in 10mL of the same media, spun down again and finally, resuspended in 1-2mL media. Cells were counted using a hemocytometer and loaded on to arrays for single-cell sequencing as described for patient tissues.

#### Immunofluorescence

Organoids were passaged into 8-well Nunc Lab-Tek II Chamber Slides (Thermo Scientific, Cat: 154453) and allowed to grow in prostate-specific media. Following seven days, the media was removed, domes were washed twice with 300uL PBS and fixed in 4% paraformaldehyde (Electron Microscopy Sciences, Cat: 15710-S) at room temperature for 20 minutes. Individual domes were washed 3x with IF Buffer (0.02% Triton + 0.05% Tween + PBS) and blocked for one hour at room temperature with 0.5% Triton X100 + 1% DMSO + 1% BSA + 5% Donkey Serum + PBS. Following the block, domes were washed once with IF Buffer and incubated overnight with monoclonal mouse anti-Lactoferrin (Abcam, Cat: ab10110, 1ug/mL), monoclonal rat anti-Uteroglobin/SCGB1A1 (R&D Systems, Cat: MAB4218-SP, 1ug/mL), polyclonal guinea pig anti-Cytokeratin 8 + 18 (Fitzgerald, Cat: 20R-CP004, 1:100), and polyclonal chicken anti-Keratin 5 (Biolegend, Cat: 905901, 1:100). Subsequently, domes were washed 3x with IF Buffer and counterstained with Alexa Fluor 488-AffiniPure Donkey Anti-Chicken IgY (IgG) (H+L) (Jackson ImmunoResearch, Cat: 703-545-155, 1:500), Donkey anti-Mouse IgG (H+L) Cross-Adsorbed Secondary Antibody, DyLight 550 (Thermo Fisher Scientific, Cat: SA5-10167, 1:500), Donkey anti-Rat IgG (H+L) Cross-Adsorbed Secondary Antibody, DyLight 680 (Thermo Fisher Scientific, Cat: SA5-10030, 1:500), and Alexa Fluor 790 AffiniPure Donkey Anti-Guinea Pig IgG (H+L) (Jackson ImmunoResearch, Cat: 706-655-148, 1:500) containing DAPI (Sigma, Cat: D9542-5MG, 1:1000). Finally, wells were washed 3x with IF Buffer for five minutes and sealed with Prolong Gold antifade mountant (Fischer Sci, Cat: P36930). Z-stack images were captured on a Leica DCF9000 GT using Leica Application System X software.

### Quantification and Statistical Analysis

#### Sequencing and Alignment

Sequencing results were returned as paired FASTQ reads and processed with FastQC^96^ for general quality checks in order to further improve our experimental protocol. Then, the paired FASTQ files were aligned against the reference genome using a STAR aligner^97^ in the dropseq workflow (https://cumulus.readthedocs.io/en/latest/drop_seq.html). The aligning pipeline output included aligned and corrected bam files, two digital gene expression (DGE) matrix text files (a raw read count matrix and a UMI-collapsed read count matrix where multiple reads that matched the same UMI would be collapsed into one single UMI count) and text-file reports of basic sample qualities such as the number of beads used in the sequencing run, total number of reads, alignment logs. For each sample, the average number of reads was 4,875,9687, and the mean read depth per barcode was 48,586. The median and average number of genes per barcode were 767 and 1079. The median and average number of UMI were 1,335 and 2,447. The mean percentage of mitochondrial content per cell was 13.65%.

#### Single-cell clustering analysis

Cells in the samples were clustered and analyzed using customized codes based on the Seurat V3.0 package on R^20^. Cells with less than 300 genes, 500 transcripts, or a mitochondrial level of 20% or greater, were filtered out as the first QC process. Then, by examining the distribution histogram of the number of genes per cell in each sample, we set the upper threshold for the number of genes per cell in each individual sample in order to filter potential doublets. A total of 22,037 cells were acquired using these thresholds. Since merging with and without integration of the samples showed no major difference in the clustering of each cell type, in the subsequent analysis of these samples we used the merged dataset without integration.

Doublets were removed by two steps: first we used DoubletFinder^98^ and a theoretical doublet rate of 5% to locate doublets in our dataset. 294 cells marked by DoubletFinder as true positive were removed from further analysis. 21,743 cells were used in the following cell clustering analysis. Then, after clustering, we removed cells expressing biomarkers from more than one major cell type (epithelial, stromal, and immune) as they were more likely to be doublets. In this step, we removed 276 cells from our dataset and the follow-up analysis, leaving 21,467 cells in total.

UMI-collapsed read counts matrices for each cell were loaded in Seurat for analysis^20^. We followed the standard workflow by using the “LogNormalize” method that normalized the gene expression for each cell by the total expression, multiplying by a scale factor 10,000 and log-transforming the results. For downstream analysis to identify different cell types, we then calculated and returned the top 2,000 most variably expressed genes among the cells before applying a linear scaling by shifting the expression of each gene in the dataset so that the mean expression across cells was 0 and the variance was 1. This way, the gene expression level could be comparable among different cells and genes. PCA was run using the previously determined most variably expressed genes for linear dimensional reduction and the first 100 principal components (PCs) were stored which accounted for 25.42% of the total variance. To determine how many PCs to use for the clustering, a JackStraw resampling method was implemented by permutation on a subset of data (1% by default) and rerunning PCA for a total of 100 replications to select the statistically significant principle component to include for the K-nearest neighbors clustering. For graph-based clustering, the first 100 PC and a resolution of 3 were selected yielding a total of 46 cell clusters. We eliminated the clustering side effect due to overclustering by constructing a cluster tree of the average expression profile in each cluster and merging clusters together based on their positions in the cluster tree. As a result, we ensured that each cluster would have at least ten unique differentially expressed genes (DEGs). Differentially expressed genes in each cluster were identified using the FindAllMarker function within Seurat package and a corresponding p-value was given by the Wilcoxon’s test followed by a Bonferroni correction. Top differentially expressed gene markers were illustrated in a stacked violin plot using a customized auxiliary function. Dot plots were generated as an alternative way of visualization using the top ten differentially expressed genes in each cluster. Top tier cell type clustering was also validated by the automated singleR annotation (**Supplemental Table 1**)

#### Cell type annotation by signature scores

In order to annotate each cell type from the previous clustering, we took the established studies and the signatures for each cell type (**Supplemental Table 2**). Treating the signature score of each cell type as a pseudogene, we evaluated the signature score for each cell in our dataset using the AddModuleScore function^20^. Each cluster in our dataset was assigned with an annotation of its cell type by top signature scores within the cluster.

#### Epithelial sub-clustering analysis and tumor cell inference

All epithelial cells were clustered using the analytical workflow described above, yielding 20 clusters. To compare the transcriptomic profiles between PCa samples and normal prostates, a previous study on normal prostate single-cell RNA-seq was downloaded and imported. Mean basal, luminal, hillock, and club signature scores were calculated for each cluster, based on the top differentially expressed genes from a previous scRNA-seq study on the normal prostate. A One-way ANOVA test was then conducted to determine if the signature score of each cluster was significantly different from the rest. We annotated the clusters with significantly upregulated basal epithelial cell (BE) signature scores as BE. Cells in clusters with high luminal epithelial (LE) signature scores could be either non-malignant luminal epithelial cells or tumor cells. The clusters with low signature scores of both BE and LE were annotated as other epithelial cells (OE). To efficiently identify tumor cells, we took the digital gene expression matrix and conducted a single set gene set enrichment analysis on GenePattern (https://gsea-msigdb.github.io/ssGSEA-gpmodule/v10/index.html) testing against the C2 gene set collection curated on MSigDB (https://www.gsea-msigdb.org/gsea/msigdb/index.jsp). Under the notion that tumor cells should have higher expression of one or more tumor markers overlapping existing prostate cancer gene sets, we projected the signatures of these prostate cancer gene sets on to our epithelial clusters and annotated tumor cell clusters as the clusters with significantly higher (p < 0.05 in one-way ANOVA test) signature scores of at least one prostate cancer gene sets.

Approximately ∼50% of prostate cancer cells from men of European ancestry harbor *TMPRSS2-ERG* fusion events, indicating high gene expression of *ERG*^99, 100^. Therefore, we hypothesized a high signature score of SETLUR PROSTATE CANCER TMPRSS2 ERG FUSION UP gene set^26^ would be a strong indicator of *ERG*+ tumor cells. All the other tumor cell clusters were then annotated as *ERG-* tumor cell clusters as they showed little to no *ERG* gene expression. All of the epithelial clusters with high luminal signature scores and high expression of luminal markers such as *KLK3, KLK2, ACPP, KRT8,* and *KRT18* were annotated as non-malignant luminal epithelial cells (non-malignant LE). Compared to non-malignant cells, tumor cells harbor more single-nucleotide variants and copy number variants, leading to distinctive patterns. To validate our tumor cell annotation, we ran InferCNV on *ERG*+ and *ERG*- tumor clusters with non-malignant LEs as reference^29^ for an estimation of copy number alterations. We classified tumor cells based on *ERG* gene expression. Then we defined patients harboring *ERG*+ tumor cells as *ERG*+ patients and the other patients as *ERG*- patients. This way, we were able to classify all the other cells based on the *ERG* status (epithelial, stromal, and immune cells) as either *ERG*+ or *ERG*-.

To determine if common functional changes were present in more than one cell type, we conducted gene set enrichment analysis (GSEA) for each cell type first and imported the significantly changed gene sets to take the intersections. Statistical significance of multi-set intersection was evaluated and visualized using the SuperExacTest package^51^.

#### Cell state analysis

Gene expression profile differences in epithelial cells between PCa sample and normal prostate samples were identified by integrating our PCa dataset with an established dataset on normal prostates^9^. We utilized the integration method based on commonly-expressed anchor genes by following the Seurat integration vignette^20^ in order to remove batch effects of samples sequenced with different technologies and possible artifacts so that the cells were comparable.

In order to better characterize the transcriptomic profile and transition of cell states among identified epithelial cells, both the tumor and paired normal samples were integrated together and separately with the epithelial cells from a normal prostate scRNA-seq dataset^9^ for *KRT5*+ and *KRT15*+ basal epithelial (BE), *KLK3*+ and *ACPP*+ luminal epithelial (LE), and *PIGR*+ and *MMP7*+ club cell population together and separately. An optimal resolution value was tested using the Clustree^101^ package. Heatmaps of DEGs were generated to validate the cell state differentiation. Compositions for each cell state were computed and compared between PCa samples and normal samples using Fisher’s exact test.

To assess the functional roles of the PCa-enriched cell states identified within the integrated dataset, we ran GSEA analysis between the PCa-enriched cell state and all the other cell states as a whole. The top 20 downregulated and upregulated gene sets were visualized in terms of gene counts and ratio for each gene set. Using the DEGs from each cell state, we generated signature gene sets for all the cell states in BE, LE, and club cells. To validate the functional implications for the PCa-enriched cell states, we conducted ssGSEA on PCa BE and club cells to compute the signature scores of the upregulated gene sets using the ssGSEA module on GenePattern (https://gsea-msigdb.github.io/ssGSEA-gpmodule/v10/index.html). Then, we computed the information coefficient (IC) and corresponding p-values followed by FDR correction to evaluate the correlation between these gene sets and cell states.

#### Pseudotime analysis

To evaluate the epithelial cell states with respect to their order in the differentiation trajectory, we conducted pseudotime analysis on all epithelial and tumor cells identified in the PCa samples. We first calculated a PAGA (partition-based graph abstraction) graph using SCANPY’s sc.tl.paga() function^102^ and then used sc.tl.draw_graph() to generate the PAGA initialized single-cell embedding of the cell types (**Supplemental Figure 4a)**. The diffusion pseudotime for each cell was calculated using SCANPY’s sc.tl.diffmap() and sc.tl.dpt() with the root cell chosen from the stem cell upregulated BE cluster and then was plotted on the PAGA initialized embedding. **(Supplemental Figure 4b)**. We then visualized the gene marker changes along the pseudotime by cell type using sc.pl.paga_path() (Supplemental Figure 4c).

Furthermore, to test whether or not the luminal-like cell state within the club cell population was more differentiated compared to other club cells, we utilized Monocle3^45^ on club cells. Monocle3 object was generated using the count matrix for all club cells and the pseudotime trajectory was computed following the standard Monocle3 workflow. The starting point of the trajectory was identified using the cell with the highest adult stem cell signature score (**Supplemental Figure 4d**) and the luminal-like club cells were highlighted using the luminal epithelial cell signature (**Supplemental Figure 4d**). Expression levels along the pseudotime trajectory for club cell markers *LTF* and *PIGR,* and luminal markers *ACPP* and *KLK3* were then plotted.

#### scRNA-seq Fusion detection

Fusion transcripts were detected using STAR-Fusion^27^ (https://github.com/STAR-Fusion/STAR-Fusion/wiki) version 1.6.0. STAR-Fusion was run from a Docker container using the following options: *--FusionInspectorvalidate, --examine_coding_effect, and – denovo_reconstruct.* Due to the low coverage of scRNA-seq samples both filtered fusion detection results and preliminary results were combined and processed, and we only filtered for potential *TMPRSS2-ERG* fusion events.

#### Signature analyses of bulk RNA-sequencing datasets

Two publicly available bulk RNA-sequencing PCa datasets were used to test the correlation between the PCa-enriched cell state signatures and AR signaling, including Prostate Adenocarcinoma (TCGA^25^, Firehose Legacy) dataset (N = 499, available at https://www.cbioportal.org/study/summary?id=prad_tcga) and SU2C/PCF Dream Team (SU2C^48^, PNAS 2019) dataset (N = 266, available at https://www.cbioportal.org/study/summary?id=prad_su2c_2019). For each dataset, mRNA expression was downloaded and normalized. Signature scores of *AR* signaling (Hallmark androgen response pathway), BE, LE, and club cell states as well as *ERG+* and *ERG-* tumor cell signature scores were computed for each sample via ssGSEA. Samples in each dataset were rank ordered by the *AR* signature scores and heatmaps were generated using the customized scripts. To test the correlation between *AR* signature scores and each cell state signature score, we computed the information coefficient and corresponding p-values followed by FDR correction to evaluate the correlation. For tumor cell signatures, we computed the correlations between the *ERG* fusion status from each dataset and the signature scores of *ERG+* and *ERG-* tumor cell gene sets we had previously generated. We rank ordered the bulk RNA-seq samples according to whether or not the *TMPRSS2-ERG* fusion was detected and plotted the *ERG*+ and *ERG*- tumor cell signature score heatmaps. Information coefficient (IC), p-values, and FDR q-values were computed.

#### Immune cell analysis

T-cell and myeloid cell populations were sub-clustered separately following a similar pipeline as described above. For T-cells, 23 PCs and a resolution of 1.5 were selected for the clustering. For myeloid cells, 27 PCs and a resolution of 1.5 were selected. Cell clusters were annotated by a dot plot showing the top ten most expressed genes in each cluster.

Monocytes, macrophages, neutrophils, and eosinophils were identified and annotated based on the automated SingleR analysis^19^. M1/M2 macrophage phenotypes, tumor associated macrophages, and two types of myeloid-derived suppressor cells were identified using documented markers from previous studies.

### Materials Availability

This study did not generate new unique reagents.

### Data and Code Availability

Processed single-cell RNA sequencing data that support this study will be deposited in the NCBI GEO database and available upon request to the corresponding author. All software algorithms used for analysis are available for download from public repositories. All code used to generate figures in the manuscript are available upon request.

## Acknowledgments

This work was in part supported by: Searle Scholars Program (A.K.S.), Beckman Young Investigator Program (A.K.S.), Sloan Fellowship in Chemistry (A.K.S.), Pew-Stewart Scholars Program for Cancer Research (A.K.S.), and the Prostate Cancer Foundation (F.W.H.).

We thank Travis Hughes and Matthew Hellmann for helpful discussions.

## Author Contributions

Conceptualization, H.S and F.W.H.; Methodology, H.S, H.N.W, P.A, M.H.W, B.W., and F.W.H.; Investigation, H.S. and J.X., Writing – Original Draft, H.S. and F.W.H.; Writing – Review & Editing, H.S., H.N.W., P.A., J.X., M.H.W, F.Y.F. M.R.C., and A.K.S., F.W.H.,; Resources, M.R.C., P.C., B.W., H.Y., A.K.S. and F.W.H.; Supervision, F.W.H.

