## Supplemental Information for "Single-cell analysis of human primary prostate cancer reveals the heterogeneity of tumor-associated epithelial cell states"

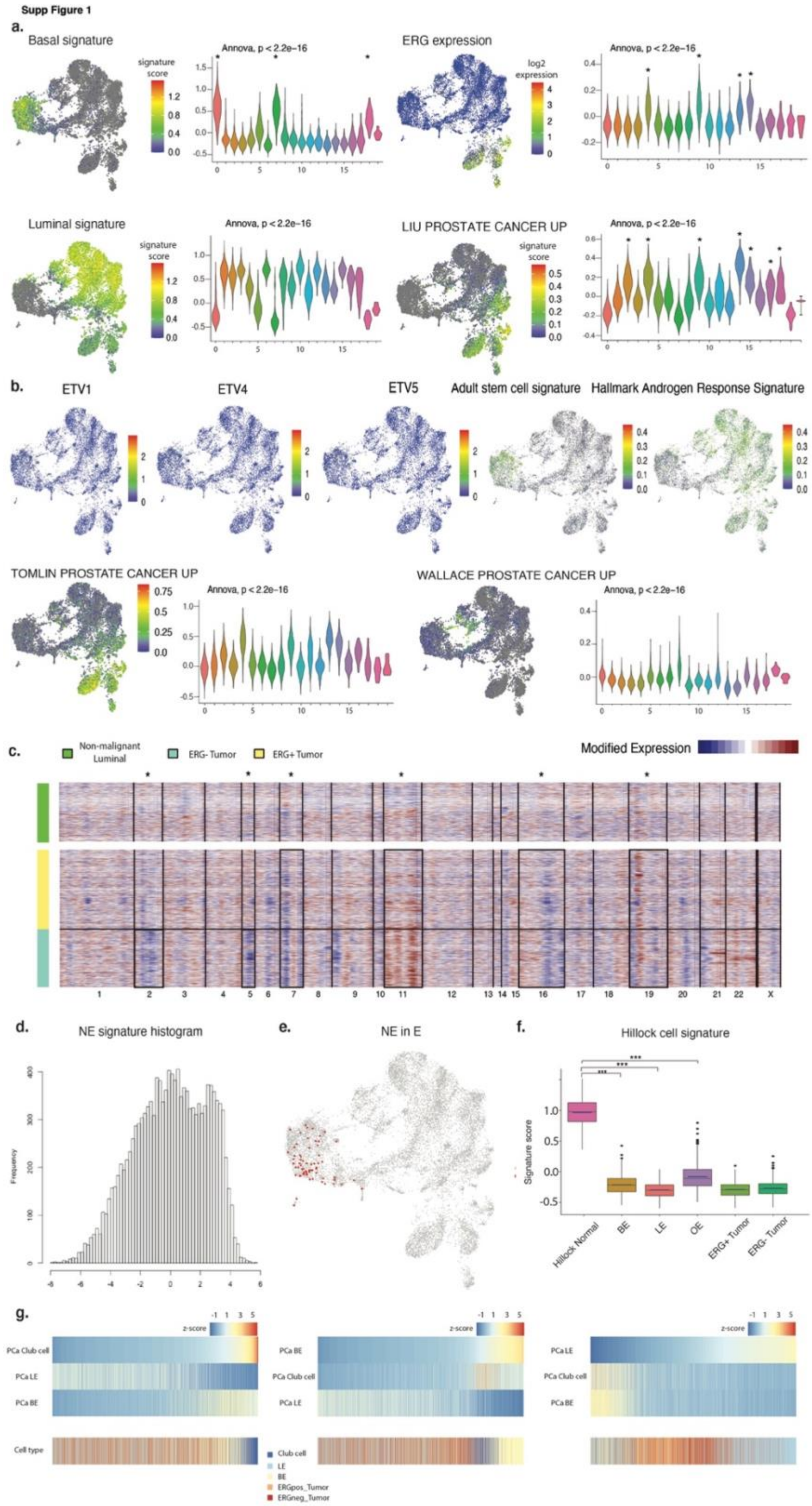

**Supplemental Figure 1. Epithelial cell type annotation and validation. a.** Left, UMAP projection of normal basal epithelial (BE), luminal epithelial (LE) cell signatures, *ERG* expression and LIU PROSTATE CANCER UP signature, with corresponding violin plots for epithelial cell clusters (p-values of one-way ANOVA test are labeled, clusters with upregulated signature scores are highlighted with asterisks). **b.** Top, Featureplots of *ETV1, ETV4, ETV5* expression, adult stem cell signature score, and Hallmark androgen response signature score. Bottom, UMAPs of Tomlin prostate cancer up and Wallace prostate cancer up tumor marker gene sets signature scores with corresponding violin plots. **c.** InferCNV result with significant copy number variations highlighted. **d.** Distribution of neuroendocrine (NE) signature score. **e.** UMAP of epithelial cells with identified NE highlighted in red. **f.** Hillock cell signature score comparison between PCa epithelial cells and hillock cells from normal samples (***: $p < 0.001$, Wilcoxon rank sum test). **g.** ssGSEA validation of PCa BE, LE, and club cell signature gene sets. z-score of ssGSEA signature scores are computed and ordered.

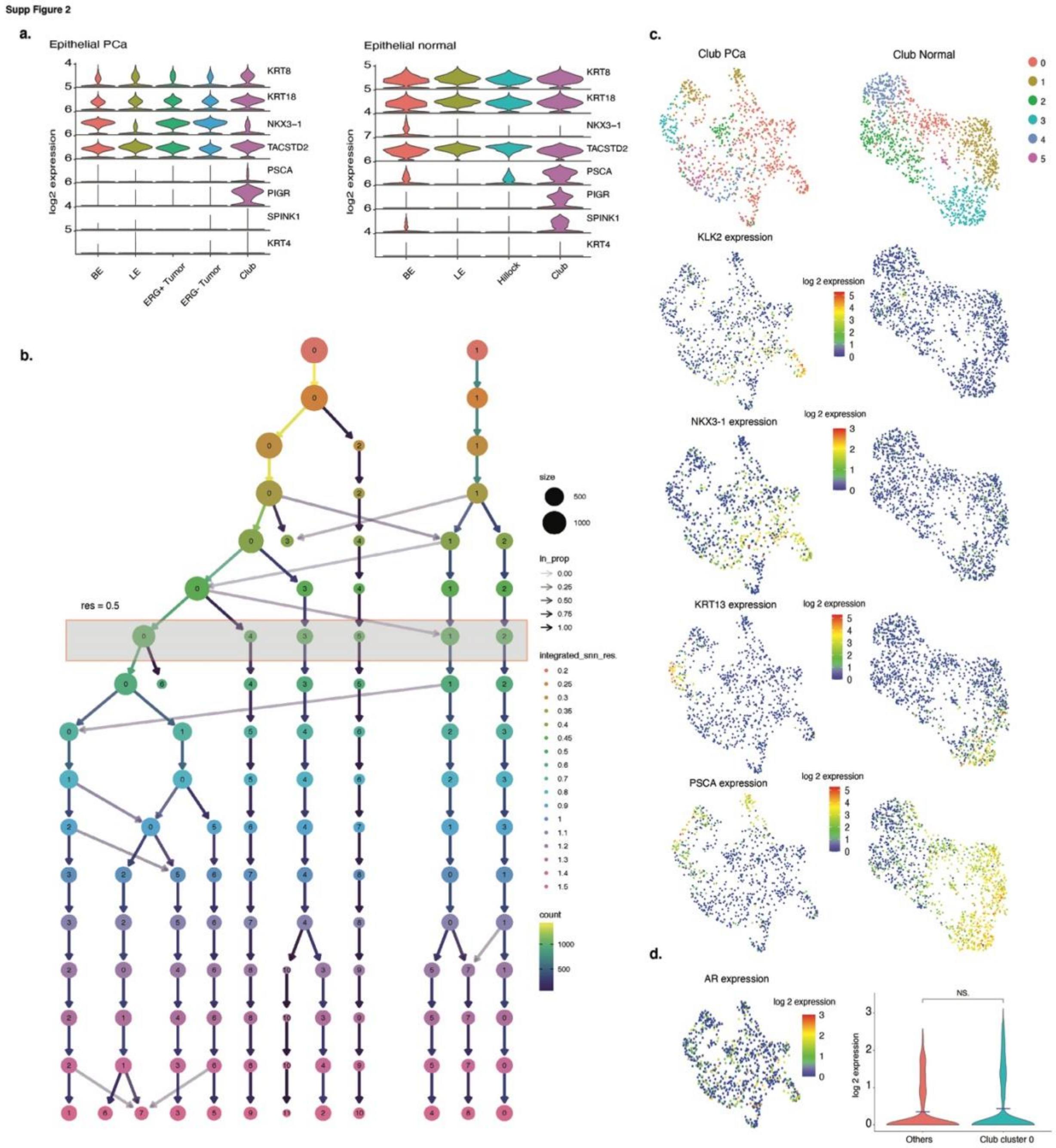


**Supplemental Figure 2. Integrated club cell analysis. a.** Stacked violin plots of LE progenitor cell markers for PCa (left) and normal (right) epithelial cells. **b.** Clustering stability tree for integrated club cells. Cluster transition is indicated by arrows and cluster sizes are indicated by marker sizes. Selected resolution for downstream analysis is labeled next to the highlighted region. **c.** Computed UMAP for club PCa and club

Normal. Featureplots show top expressed genes in different cell states. **d.** Comparison of *AR* expression between club cell cluster 0 and other club cells in the PCa samples (NS: not significant, Wilcoxon rank sum test).

Supp Figure 3

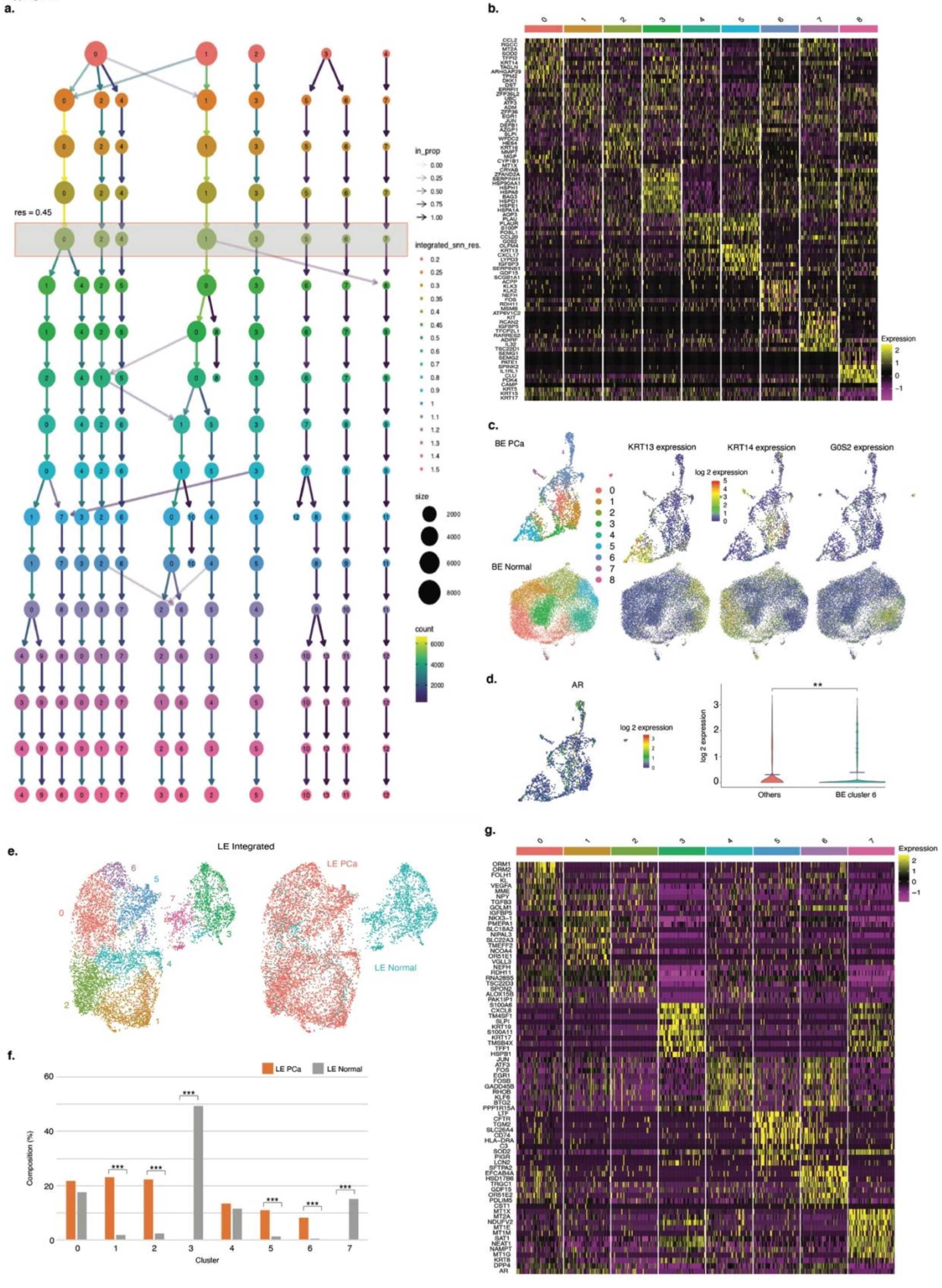

**Supplemental Figure 3. Integrated BE and LE analysis. a.** Clustering stability tree for integrated BE. Cluster transition is indicated by arrows and cluster sizes are indicated by marker sizes. Selected resolution for downstream analysis is labeled next to the highlighted region. **b.** Heatmap of the top 10 differentially expressed genes (DEGs) for each BE cell state. **c.** Computed UMAP for BE PCa and BE Normal. Featureplots show top expressed genes in different cell states. **d.** Comparison of *AR* expression between BE cell cluster 6 and other BE in the PCa samples (**: $p < 0.01$, Wilcoxon rank sum test). **e.** UMAP of integrated LE labeled by cell states (left) or samples type (LE PCa and LE Normal) (right). **f.** Cell composition comparison between LE PCa and LE Normal. **g.** Heatmap of the top 10 DEGs for each LE cell states.

### Supp Figure 4

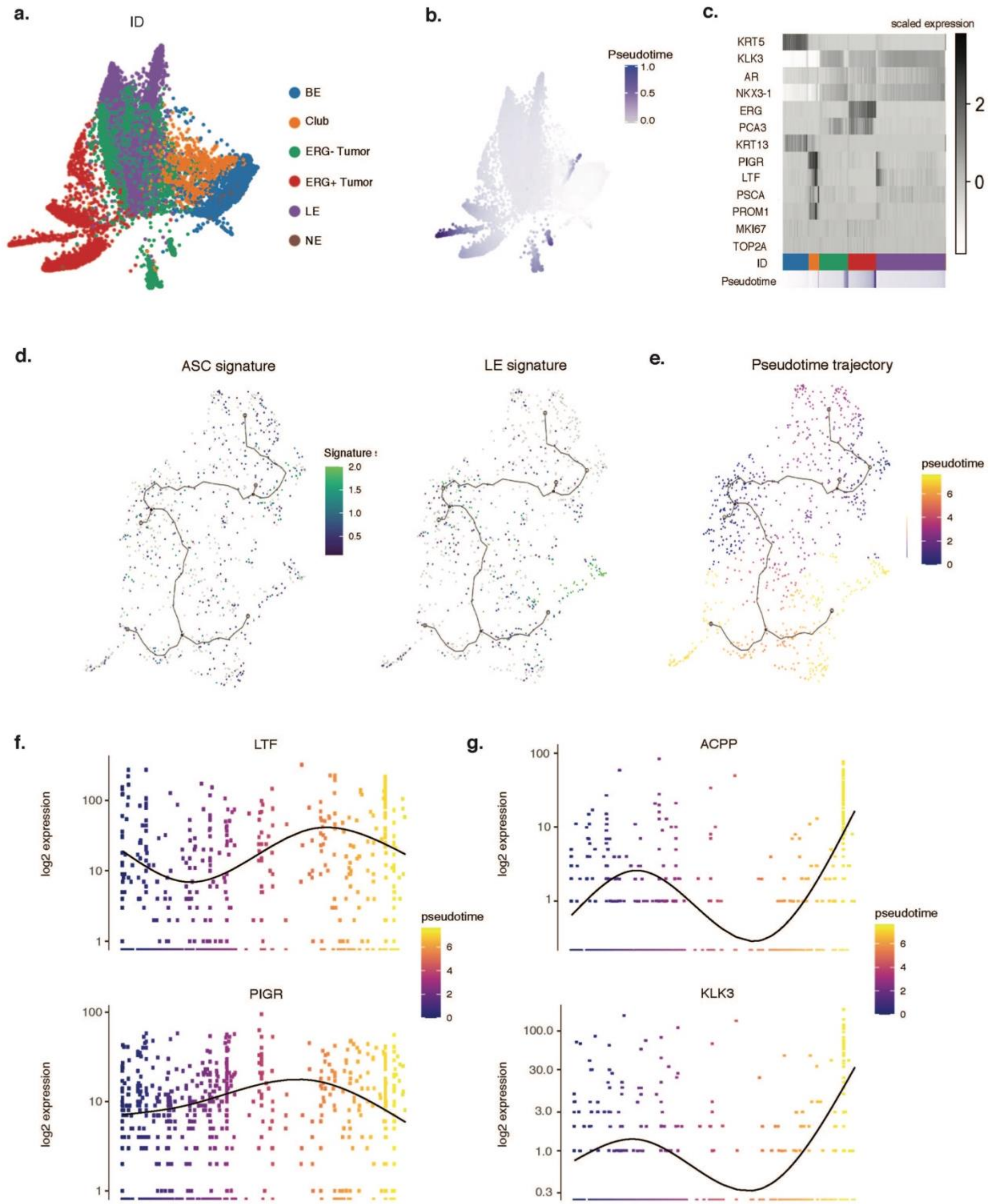

**Supplemental Figure 4. Pseudotime analysis on PCa epithelial cells. a.** Projection of all epithelial cells in the pseudotime plane. **b.** Pseudotime heatmap for the epithelial cells. Computed trajectory is shown in the color bar. **c.** Expression changes along the pseudotime for BE, LE, tumor, club cell and proliferation markers. **d.** monocle3 reduction UMAP of PCa club cells for adult stem cell (ASC) and LE signature scores. **e.** Pseudotime trajectory for all club cells. **f.** Expression of club cell markers *LTF* and *PIGR* along the pseudotime trajectory. **g.** Expression of LE markers *ACPP* and *KLK3* along the pseudotime trajectory.

**Supp Figure 5**

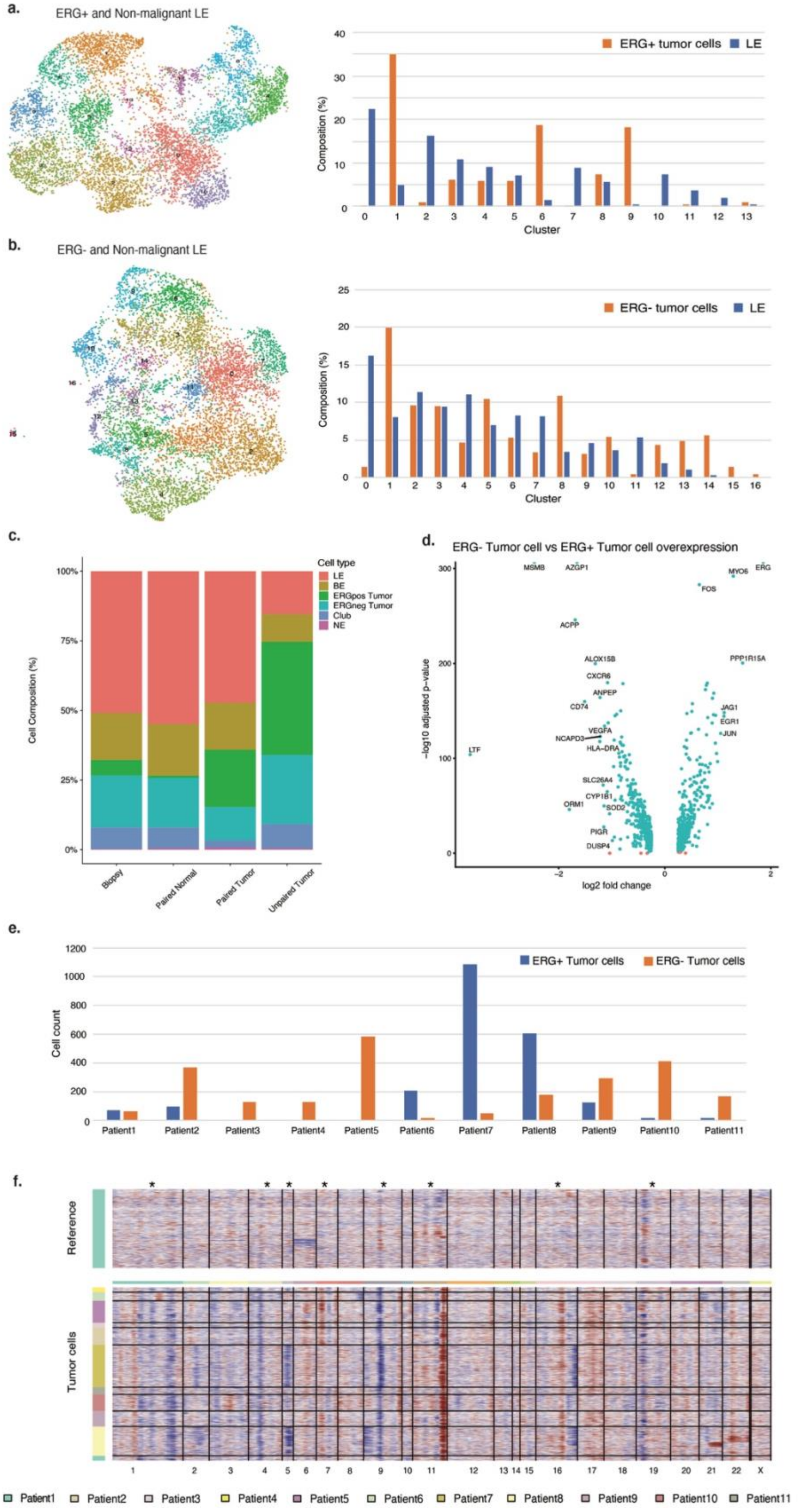

**Supplemental Figure 5. Tumor cell analysis. a.** Left, UMAP of integrated *ERG*+ tumor cells and LE. Right, cluster composition comparison in grouped bar charts. **b.** Left, UMAP of integrated *ERG*- tumor cells and LE. Right, cluster composition comparison in grouped bar charts. **c.** Stacked bar chart of epithelial cells in each sample type. Composition is normalized to 100%. **d.** Volcano plots between *ERG*+ and *ERG*- tumor cells with the top 20 most overexpressed genes labeled. **e.** Group bar charts of both *ERG*+ and *ERG*- tumor cell counts in each patient. **f.** InferCNV validation of tumor cells by patients using non-tumor epithelial cells as reference.

**Supp Figure 6**

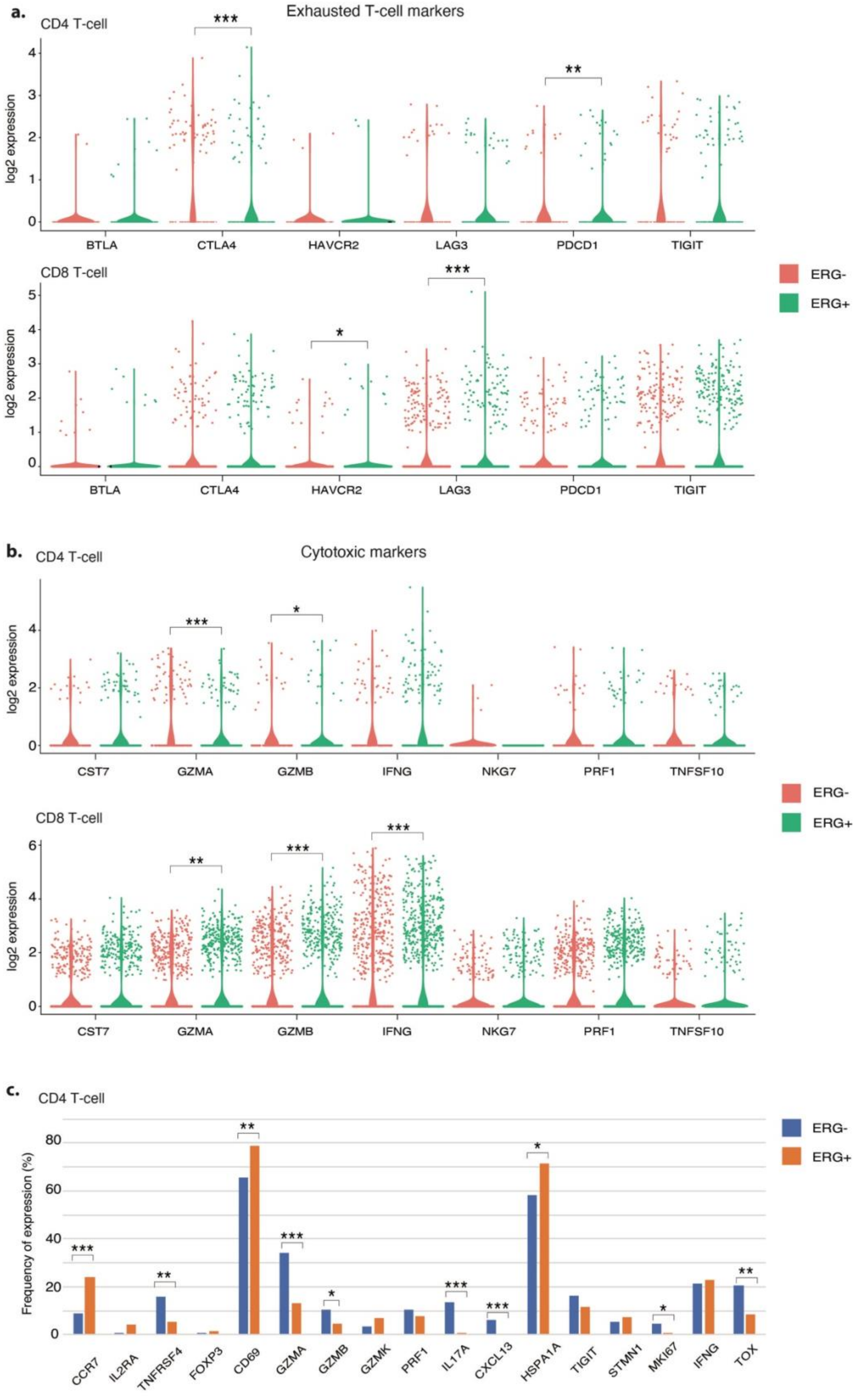

**Supplemental Figure 6. *ERG*-associated CD4 and CD8 T-cell analysis. a.** Grouped violin plots of exhausted T-cell markers for CD4 and CD8 T-cells classified by the *ERG* status. **b.** Grouped violin plots of cytotoxic markers for CD4 and CD8 T-cells classified by the *ERG* status. Statistical significance is labeled above (*: $p < 0.05$; **: $p < 0.01$, ***: $p < 0.001$, no label: not significant, Wilcoxon rank sum test). **c.** Frequency of expression for CD4 T-cell subtype markers between the two CD4 T-cells classified by the *ERG* status (*: $q < 0.05$; **: $q < 0.01$, ***: $q < 0.001$, no label: not significant, FDR).

Supp Figure 7

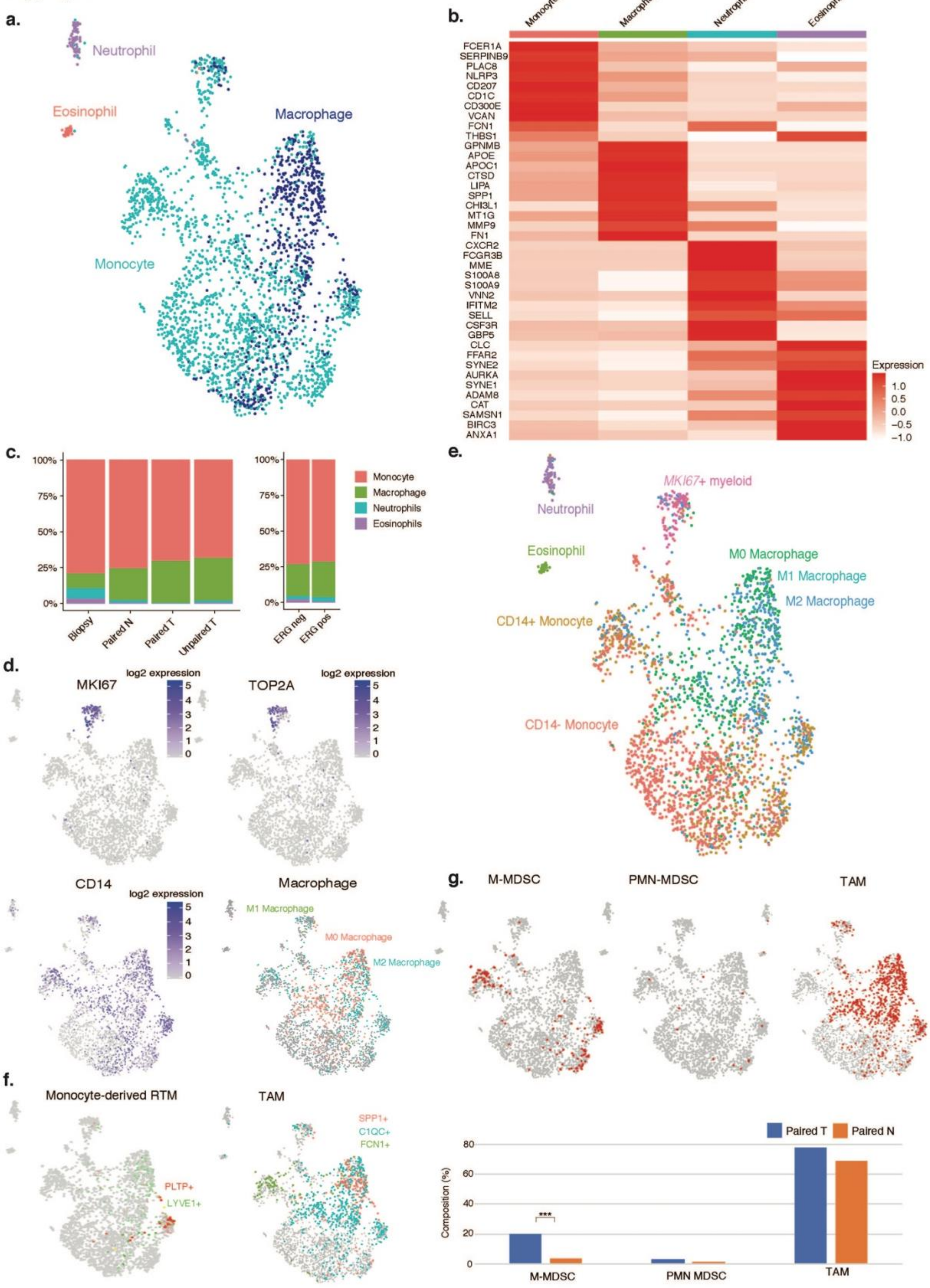

**Supplemental Figure 7. Myeloid cell analysis. a.** UMAP of myeloid cell population with SingleR automated annotation. **b.** Heatmap of the top 10 DEGs in each myeloid cell type. **c.** Myeloid cell composition comparison by each sample type (left) and by *ERG* status (right). **d.** Featureplots of *MKI67*, *TOP2A,* and *CD14*; distribution of M0, M1, and M2 macrophages in the myeloid cell UMAP. **e.** UMAP of myeloid cells annotated by detailed monocyte and macrophage phenotypes. **f.** Featureplots of monocyte-derived resident tissue macrophages (RTM) markers and tumor associated macrophage (TAM) markers. **g.** Top, identification of two myeloid-derived suppressor cell (MDSC) phenotypes and TAMs within myeloid cells. Bottom, composition comparison between paired tumor and paired normal samples (***: $p < 0.001$, FET).

Supp Figure 8

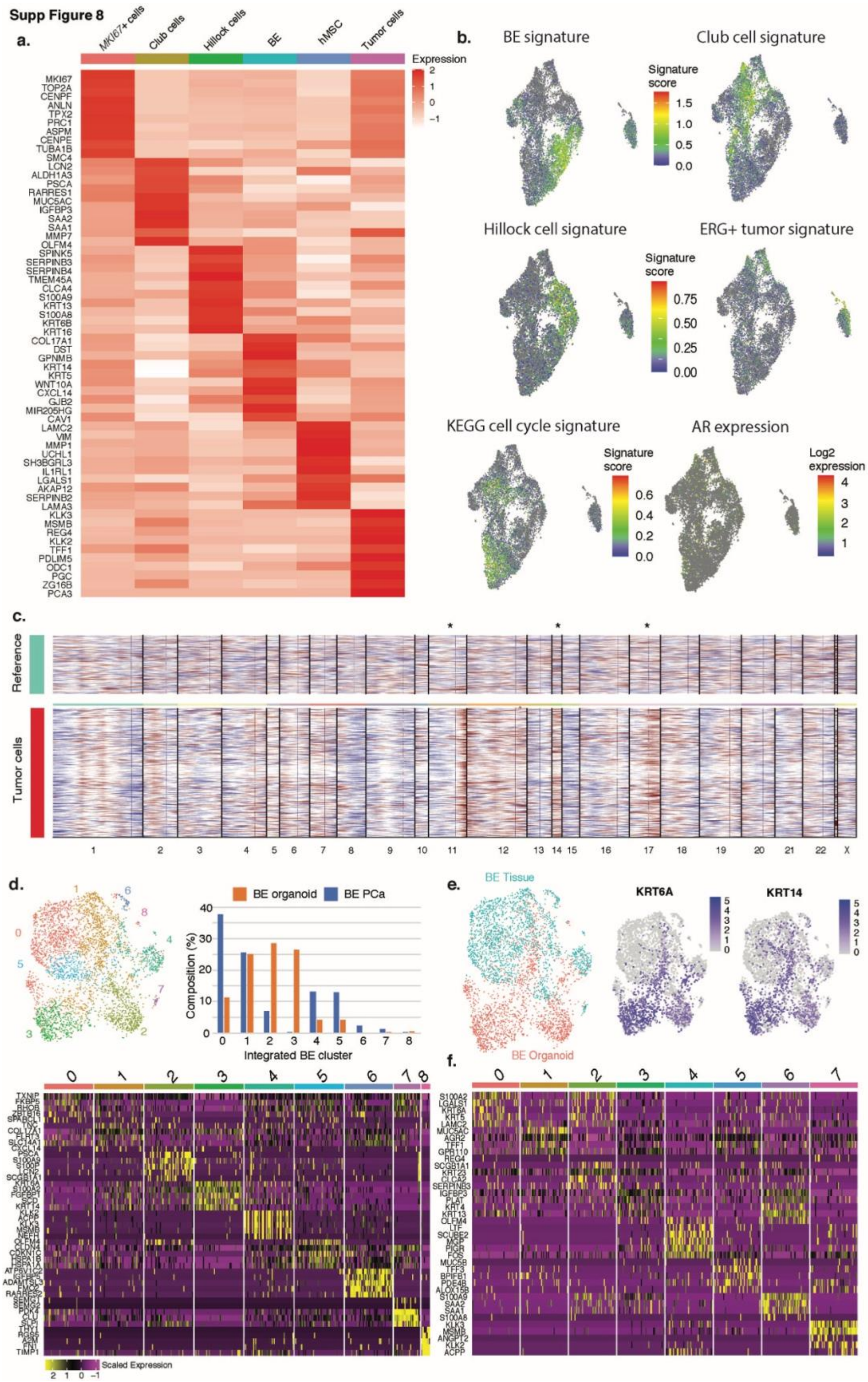

**Supplemental Figure 8. Analysis on early-passage (P0-P3) organoid samples.** **a.** Heatmap of the top 10 DEGs for each organoid cell type. **b.** UMAP of different tumor tissue cell type signature scores and *AR* expression Featureplot in the organoid samples. **c.** InferCNV validation using non-tumor cells as reference and tumor cells as observation. Significant copy number variations are highlighted. **d.** Top, UMAP of integrated BE dataset of tumor tissue and organoid samples and cell composition comparison grouped bar charts. Bottom, heatmap of the top 10 DEGs in the integrated BE clusters. **e.** Left, UMAP of the integrated BE cells labeled by sample type. Right, Featureplots of organoid specific BE cell state markers *KRT6A* and *KRT14*. **f.** Heatmap of the top 10 DEGs in the integrated club cell clusters.

**Supplemental Table 1. Pathological summary for all 11 PCa patients studied in this analysis and top-tier cell type annotation validation by SingleR.**

**Supplemental Table 2. Signature gene sets for each major cell type.**

**Supplemental Table 3. ssGSEA results for all 20 epithelial cell clusters.**

**Supplemental Table 4. GSEA results for PCa-enriched club and BE cell states compared to other club cells and BE within PCa samples.**

**Supplemental Table 5. Differentially expressed gene list for three major epithelial cell types between paired tumor and paired normal samples.**

**Supplemental Table 6. T-cell subtype marker comparison between *ERG*+ and *ERG*- patients.**

**Supplemental Table 7. SingleR automated annotation results of myeloid cells and stromal cells with corresponding statistical significance.**
